# An Unprecedented [Cu-4Fe-4S] Cluster in a Methanotroph Acetol Dehydrogenase

**DOI:** 10.64898/2026.06.11.731722

**Authors:** C. Liu, E. Andreeva, A. Pol, T.R.M. Barends, X. Ye, K. Sengupta, S. DeBeer, G.E. Cutsail, H.J.M. Op den Camp, L.J. Daumann, W. Versantvoort

## Abstract

Biocatalytic metal-containing clusters are nanometer-sized chemical reactors that enable enzymes to perform chemistry far beyond what would be possible with amino acids alone ^1,2^. These clusters display a remarkable variability^3^ and elucidating their exact mechanisms is often difficult to determine, requiring input from a multitude of techniques ranging from spectroscopic to structural and theoretical methods. Thus, for unknown clusters, it is essential to accumulate, collate, and interpret as much information as possible from every relevant technique available. Here we report the discovery and in-depth, interdisciplinary characterization of an entirely novel, [Cu-4Fe-4S] cluster in the protein acetol dehydrogenase (AceDH). AceDH, isolated directly from *Methylacidiphilum fumariolicum* SolV cells, catalyzed the oxidation of acetol to methylglyoxal, proving its role in the 2-propanol/acetone metabolism of methanotrophs^4–6^. Structural, spectroscopic and electrochemical analyses reveal the [Cu-4Fe-4S] cluster has a unique three-dimensional- and electronic structure involving electronic coupling between the copper and one of the iron atoms, likely contributing to its high, +275 mV, redox potential. The binding site for the novel cluster is composed of two protein subunits and involves a novel motif. These findings expand the known repertoire of biological metal cofactors and provide insight into how heterometallic clusters are adapted for biological processes.

---

Recently, the metabolic versatility of the thermoacidophilic verrucomicrobial methanotrophs^7^ was expanded with the ability to catabolize C3 compounds, for which a metabolic pathway was inferred through genomics and transcriptomics^4,5,8^. Here we purified the enzyme acetol dehydrogenase (AceDH) directly from the bacterium *Methylacidiphilum fumariolicum* SolV growing on 2-propanol/acetone^5^. AceDH is able to oxidize acetol to the toxic compound methylglyoxal by AceDH with a *k_cat_* of 0.69 ± 0.01 s^-1^, and a *K_M_* of 2.3 ± 0.2 µM acetol (**Figure 1A**, Supplementary Text, **Extended Data Figure 1**), thereby converting the product of acetone oxidation by pMMO3^4^. Protein sequencing indicated AceDH to consist of the gene products of the *gmcA* and *gmcB* genes. The product of the *gmcB* gene, which we call the AceDH “β” subunit, contains an N-terminal Tat signal peptide, suggesting involvement in transporting the AceDH protein to the periplasm (Supplementary Information). The *gmcA* gene was found to encode a 526 amino acid-long member of the glucose-methanol-choline (GMC) family of flavoproteins ^9^ (the AceDH “α” subunit, Supplementary Information). However, the UV-Vis spectrum of AceDH purified from *M. fumariolicum* SolV growing on acetone does not show the peaks at around 370 and 450 nm typical of flavin-containing proteins. Instead, the protein displays a very broad absorption with a maximum at 400 nm and shoulders on either side at 460 nm and 365 nm (**Extended Data Figure 1**). The crystal structure of the AceDH protein was determined to 1.8 Å resolution (**Extended Data Table 1**, Supplementary Information). In the crystal, the protein forms αβ heterodimers (**Figure 1A**) in which the α protomer indeed adopts the typical flavin-binding fold of a GMC oxidoreductase ^11,12^. Within the α-subunit, the apparent active site contains a covalently attached flavin adenine dinucleotide (FAD) buried inside the protein (**Extended Data Figure 1G**), only connected to the bulk solvent *via* a 13 Å long tunnel (**Figure 1B**). However, the maps further showed an electron-dense moiety in a cavity ∼11 Å distance from the flavin, towards the interface between the α- and the β protomer and close to the heterodimer’s surface. This cavity was lined with cysteine residues (Cys187 α, Cys190 α, Cys193 α, and Cys197 α), which sequence homology and Colabfold structure prediction identified as a potential iron-sulfur-cluster binding motif (**C**DQ**C**GF**C**FQG**C**T) with the typical P at the last position missing. Indeed, a [4Fe-4S] cluster was found to explain most but not all of the remaining density, as it left a very electron dense feature unaccounted for. The height of this feature suggested a metal ion in this position, and various metal ions were tried in refinement, including iron, copper, and nickel, but no clear identification could be made. An X-ray fluorescence spectrum collected from a crystal (**Figure 1C**) clearly showed the presence of copper. Diffraction data were then measured using X-rays just above the iron- and copper K absorption edges, and anomalous difference density maps finally confirmed this additional atom to be a copper atom (**Figure 1D**). Moreover, B-factor refinement suggests the copper atom to be present at the same (unity) occupancy as the rest of the atoms in the cluster. Thus, the protein contains an unprecedented [Cu-4Fe-4S] cluster. This is also in line with ICP-MS measurements, which showed both iron and copper to be present in a 3:1 and 2:1 metal to protein ratio, respectively, whereas calcium, zinc, and nickel were only detected in minor amounts, with metal to protein ratios of 0.48:1, 0.01:1 and 0.001:1, respectively.

**Figure 1.**
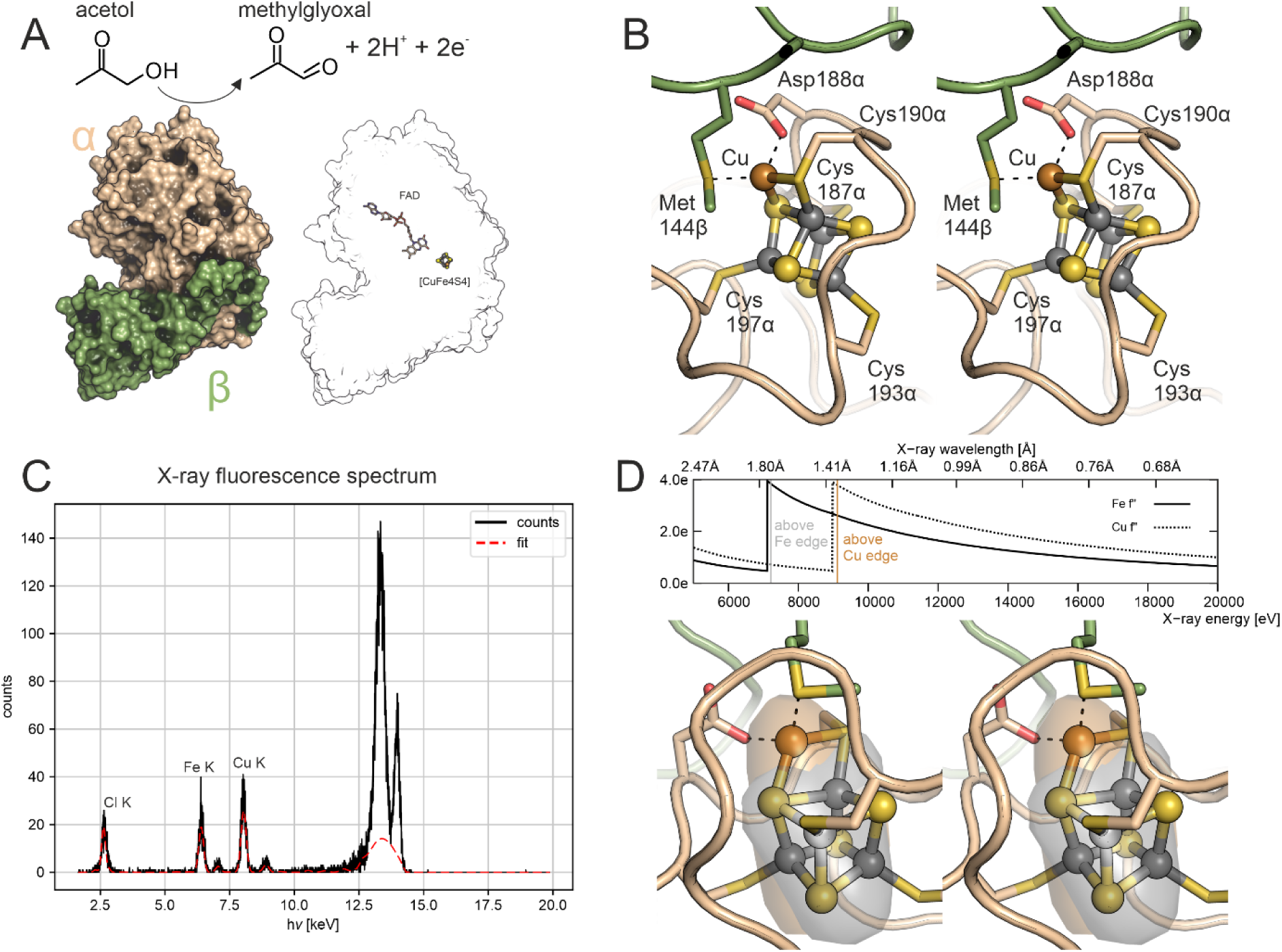
**A:** Reaction catalysed by and crystal structure of SolV AceDH. The α-subunit is shown in beige and the β-subunit in green. The panel on the right shows the position of the cofactors in the heterodimer. **B:** Stereofigure, showing the novel [Cu-4Fe-4S] cluster and its surroundings; the colouring scheme is the same as in panels B and C. The novel cluster is shown as balls and sticks with the iron atoms shown in grey, the copper atom in brown and the sulfur atoms in yellow. **C.** X-ray fluorescence spectrum from an AceDH crystal. A strong Cu K fluorescence peak is visible next to the iron K peak. **D**. Top: Wavelengths used for anomalous diffraction data collection. The f” component for the anomalous scattering is shown for iron (solid black line) and copper (dashed black line). The vertical lines show the wavelengths used to collect anomalous diffraction data sets. The data collected just above the iron edge (grey line) will show strong anomalous signal from any iron atoms, but not from a copper atom. The data collected above just above the copper edge (brown line), however, will show strong anomalous signal for both iron- and copper atoms. Bottom: stereofigure showing the anomalous difference density maps calculated from data just above the iron- and copper absorption edges, overlayed onto the structure and contoured at 5.0 σ. The “iron” map (grey surface) covers all the iron atoms but not the copper atom, whereas the “copper” map (brown surface) covers all the metal ions in the cluster.

The novel [Cu-4Fe-4S] cluster contains a four-iron-, four-sulfur-atom core in the usual shape of a cuboidal [4Fe-4S] cluster, *i.e.* with bent rhombuses forming the sides of a distorted cube (**Figure 1B, Extended Data Figure 2**). The copper is bound to the Cys190 α SG atom and a sulfur atom of the [4Fe-4S] core, forming an additional rhombus, and making the latter S atom a µ_4_ sulfur atom not seen before in iron-sulfur cluster biology. Moreover, the SG atom of Cys190 α forms a rare µ_2_ bridge between the copper and one of the irons. The copper is further bound to the SD atom of Met144β from the β protomer, and, unusually for copper, the OD1 atom of Asp188α. While the addition of a fifth rhombus to the [4Fe-4S] core may at first seem to result in a counterintuitive geometry, superimposing the novel [Cu-4Fe-4S] cluster onto other biological metal clusters shows that the distances and angles in the novel cluster are chemically plausible (**Extended Data Figure 2**). To gain further insight into the redox chemistry of this unusual cluster, we turned to spectroscopic methods.

As noted above, the UV-Vis spectrum (**Figure 2A**) of SolV AceDH shows a broad signal in the as-isolated (As is) state, with a peak around 400 nm. Addition of potassium hexacyanoferrate to the as-isolated protein did not result in significant changes in the spectrum, suggesting that as-isolated AceDH is in the oxidized state. Addition of 1 equiv. of the reducing agent ascorbate in the presence of the electron shuttle phenazine methosulfate (PMS) under anaerobic conditions showed a decrease in absorbance at 460 nm. The [as-isolated] minus [ascorbate-reduced] difference spectrum (**Figure 2A, inset**) shows peaks at 345 and 463 nm. Subsequent addition of 1 equiv. of acetol showed a further decrease in absorbance, where the [ascorbate-reduced] minus [acetol-reduced] difference spectrum (**Figure 2A, inset)** showed a peak with a maximum at 456 nm and a shoulder at 485 nm. An optical redox titration was performed, plotting the absorbance at 460 nm as a function of applied potential (vs SHE; **Figure 2B**). Two distinct redox transitions could be observed at +275 and +58 mV, respectively. A global Nernst fit was performed, extracting the components of the difference spectra corresponding to the two transitions. The +275 mV transition’s spectral component (**Figure 2B**) shows peaks at 465 nm and at ∼350 nm, which we tentatively assigned to a redox state change of the novel copper-iron-sulfur cluster. The spectral component associated with the +58 mV transition (**Figure 2C**) showed peaks at 485 nm, 456 nm and 393 nm, which we tentatively ascribed to a redox transition of the FAD cofactor. Despite the observation that the transition at +58 mV in **Figure 2C** was best fitted with a one-electron redox process, no negative difference spectrum signals above 450 nm were observed. If our assignment of this lower-potential transition to the FAD is correct, this indicates that no stable semiquinone FAD state was formed in the titrations. To corroborate our assignments and to gain further insight into the copper- and iron oxidation states of the novel cluster, we turned to electron paramagnetic resonance (EPR) spectroscopy.

**Figure 2.**
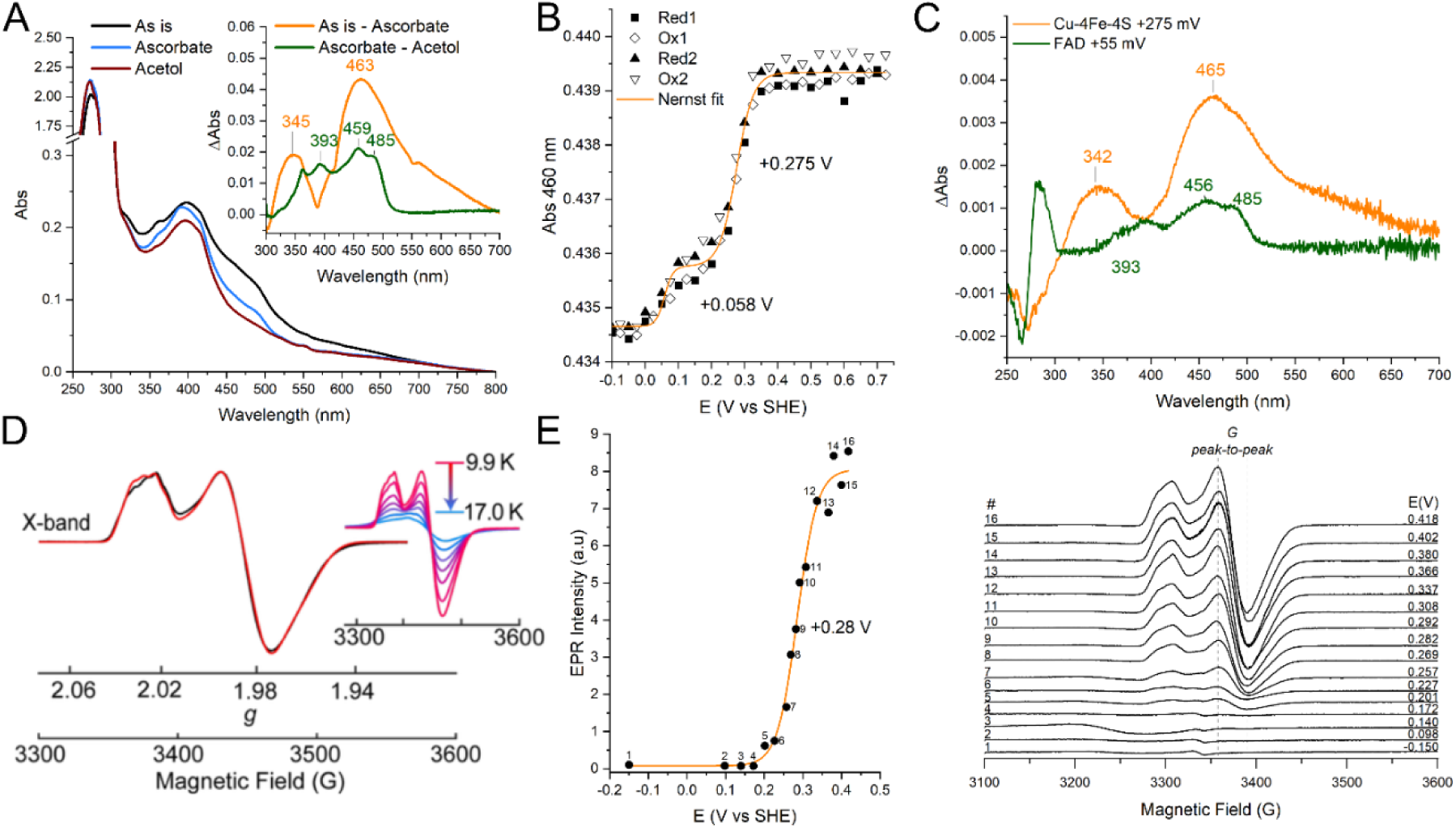
**A:** UV-Vis spectra of SolV acetol dehydrogenase. Anaerobic as isolated spectrum (Black), after addition of PMS & ascorbate (Blue) and after subsequent addition of acetol (Red). Inset: difference spectra of anaerobic as isolated minus PMS, ascorbate (Orange) and acetol, PMS & ascorbate minus PMS & ascorbate (Green). **B:** Optical redox titration of SolV AceDH, absorbance at 460 nm as a function of applied potential. AceDH was subjected to 2 reduction (closed symbols) and 2 oxidation (open symbols) cycles. The line represents the best fit to the combined reduction and oxidation data, which is a sum of 2 *n* = 1 Nernst curves. **C:** Components extracted by a global Nernst fit, corresponding to the oxidized minus reduced difference spectrum of the [Cu-4Fe-4S] cluster at +0.275 V (Orange) and to the oxidized minus reduced difference spectrum of the FAD cofactor at +0.05 V (Green). **D** X-band EPR of SolV AceDH (black) and simulation (red) with g = [2.0273, 1.9841, 1.9672], *A*_1_(Cu) = 22.7 MHz and *A*_1_(^1^H) = 21.7 MHz (complete simulation parameters given in Extended Data Table S2) **E:** EPR redox titration of SolV AceDH. Left: EPR intensity as peak-to-peak distance plotted as a function of the applied potential vs SHE. Line shows a *n* = 1 Nernst fit resulting in a reduction potential of +0.28 V. Right: Individual scans at the different redox potentials show the disappearance of the [Cu-4Fe-4S] cluster signal upon reduction. Additional EPR data is found in Extended Data Figure 4 and Spin Hamiltonian parameters in Extended Data Table S2.

The X-band (∼9.5 GHz) continuous-wave (CW) EPR spectrum of as-isolated SolV AceDH exhibits a relatively sharp *S* = 1/2 signal near g ∼ 2 that is observed only at low temperatures (<20 K, **Figure 2D**). The spectrum is approximately axial in its appearance, and simulation resolves a slight rhombic splitting; **g** = [*g*_1_, *g*_2_, *g*_3_] = [2.027, 1.984, 1.967]. Importantly, the EPR spectra give insight into the electronic state of the copper part of the cluster; along *g*_∥_ (*g*_1_), a small 6.0 to 6.7 G (16.8 to 18.8 MHz) hyperfine splitting pattern from the ^63,65^Cu nuclei is observed in the X-band EPR spectrum (**Figure 2D**), which are better resolved in the second derivative EPR spectrum (**Extended Data Figure 3**). This small coupling, along with the absence of a Cu^2+^-signal at all temperatures investigated, suggests a closed-shell Cu^+^ center^13,14^.

The determined *g*-values have a smaller shift from the free electron value (*g_e_* = 2.0023…) than what is typically observed for FeS clusters, however, the relative ordering of the *g*-values to *g_e_* and the average *g*-value (*g_av_* = 1.993 < *g_e_*) are similar to those of [4Fe-4S]^+^ clusters^15–17^. If our assignment of the +275 mV transition in the optical spectra to a redox change in the copper-iron-sulfur cluster is correct, these *g*-values are unexpected; such a high reduction potential would suggest that the novel cluster is homologous to high-potential [4Fe-4S] clusters (“Hi-PIP clusters”), and for those it has been shown empirically that *g_av_* > *g_e_* and indeed that all g-values are typically greater than *g_e_*^15^. One exception to this generalization is the Hi-PIP cluster of CoM-Hdr which exhibits very small g-shifts (g = [2.013, 1.991, 1.938]) and a *g_av_* ∼ 1.981, due in part to its perturbed FeS structure and the assigned five-coordinate iron center^18^. We therefore performed an EPR redox titration in the range of +100 to +450 mV vs SHE. The EPR signal of the [Cu-4Fe-4S] cluster is not observed at the low starting potential; only a weak, narrow radical-like signal near *g_e_* is seen (**Figure 2E**). The peak-to-peak intensity of the *g*_⊥_ feature was monitored as a function of the applied potential (**Figure 2E**) and a reduction potential of +280 mV was extracted for the [Cu-4Fe-4S] cluster cofactor (*n*=1 Nernst equation). As the protein is EPR active in the oxidized state and EPR-silent in the reduced state, the [4Fe-4S] core of the novel cluster likely switches between +2 and +3 redox states, similar to HiPIPs. The strong agreement of the redox potentials at which the transitions in the *S* = 1/2 EPR signal occur, along with the features observed at 465 nm and around 350 nm in the UV/Vis spectra, supports the identification of the copper-iron-sulfur cluster rather than FAD. Furthermore, no Cu^2+^-signals were observed throughout the whole potential range, indicating the copper remained in a Cu^+^ state.

X-ray absorption spectroscopy provides another powerful means to interrogate the iron and copper centers in this system. The Fe K-edge X-ray absorption near edge spectroscopy (XANES) spectra of the as-isolated AceDH protein exhibit a pre-edge feature at 7113 eV, followed by a rising edge at 7118 eV (**Figure 3A**). This rising edge is shifted to higher energy compared to the [4Fe-4S] cluster in the +1 oxidation state, such as the resting NifH protein^19^, indicating a more oxidized iron center in the novel cluster (**Extended Data Figure 4**). Taken together with the above spectroelectrochemical characterizations, these data indicate a more oxidized [4Fe-4S] cluster; within the framework of classical Fe–S redox assignments. In parallel, the Cu K-edge XANES spectra (Figure 3C), together with the absence of a copper(II) hyperfine interaction in the EPR spectrum, are consistent with a copper(I) center, more specifically, a four-coordinate copper(I) center (as indeed seen in the crystal structure). This is consistent with other four-coordinated copper(I) sites in complexes and proteins.^20,21^ In addition to the edge features, a weak transition was observed at approximately 8980.5 eV, which is attributed to a charge-transfer feature, and likely arises from metal-to-metal charge transfer. This again is consistent with the short distance (2.5 Å) of the copper center to the nearest iron center observed in the crystal structure (**Figures 3C and 1D**).

**Figure 3.**
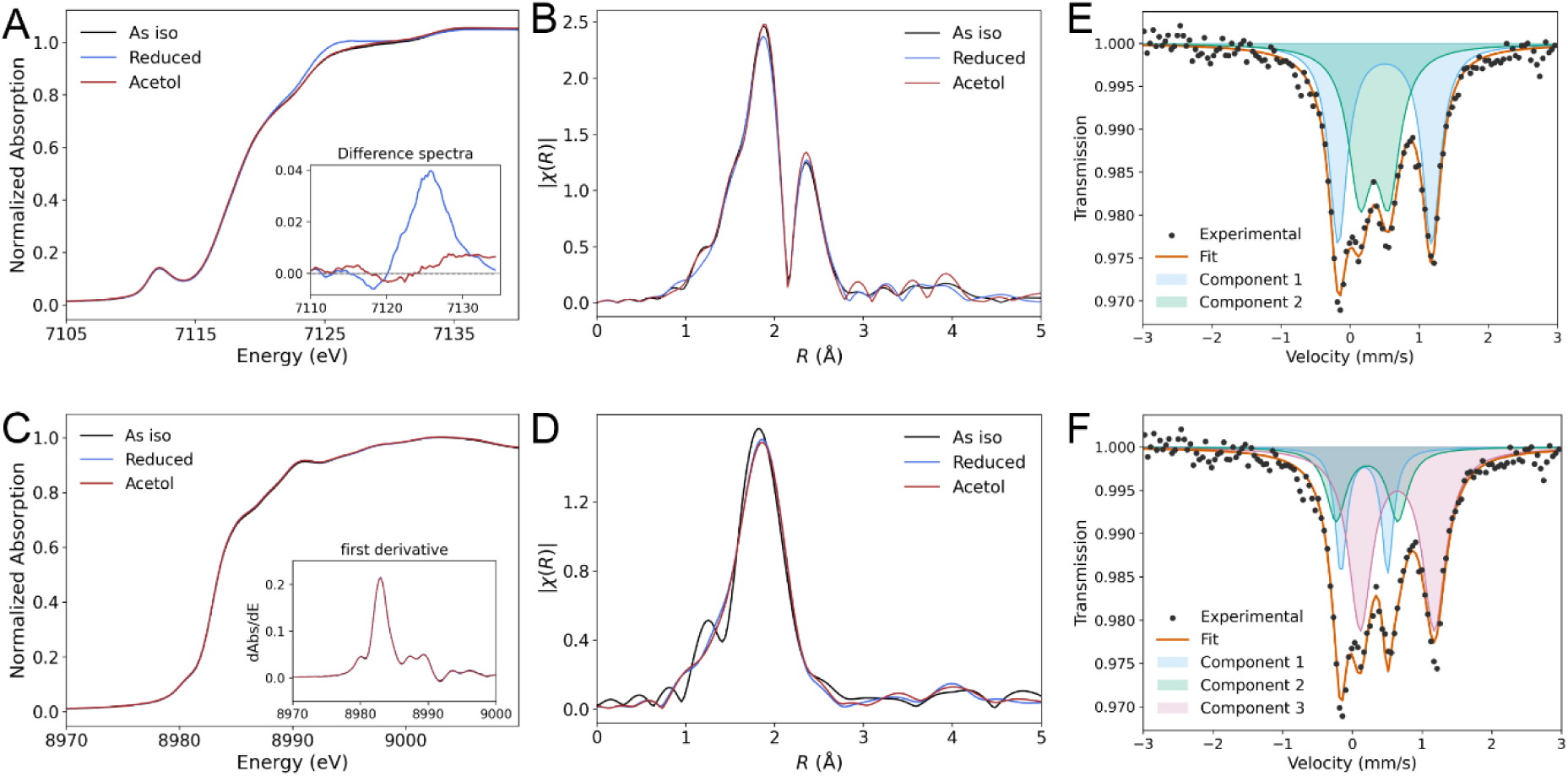
**A** Fe K-edge XANES of AceDH with the inset showing the difference spectra upon reduction and acetol adsorption, and **B** Fourier-transformed k^3^-weighted EXAFS spectra in R-space. **C** Cu K-edge XANES of the AceDH protein with the inset figure of first derivative of the spectra, and **D** Fourier-transformed EXAFS spectra in R-space. Additional EXAFS Spectra and Fits in **Extended Data Figure 4** and **Extended Data Table 2**. **E, F** Mössbauer spectrum (experimental and fitting) at 7 K (no field) of ^57^Fe enriched as isolated AceDH protein, representative fits in either a 2:2 (E) or a 1:1:2 (F) intensity ratios of the Fe doublets. Fitting results in Extended Data Table 3.

Extended X-ray absorption fine structure (EXAFS) analysis at both the Cu and Fe K-edges results in Fourier transform spectra that are dominated by features arising from first-shell Cu-S and Fe-S scattering, respectively. These primary features are complemented by weaker, longer-range contributions arising from Cu-Fe and Fe-Fe scattering pathways, in Cu and Fe FT EXAFS, respectively (**Figures 3B** and **D**, **Extended Data Table 2**). The Fe-Fe contribution can be further deconvoluted into two distinct vectors, with a shorter distance at approximately 2.5 Å and a longer distance at approximately 2.7 Å (**Extended Data Table 2**). The Fe EXAFS analysis is fully consistent with the presence of a [4Fe-4S] core in the novel cluster, and the interatomic distances obtained from fitting are in excellent agreement with those determined crystallographically (**Extended Data Table 2, Extended Data Figure 2**). Notably, Fe-Cu scattering from Fe perspective was not included as an independent path because it is obscured by overlapping Fe–Fe contributions at similar distances, making it strongly correlated with and not uniquely distinguishable from the dominant Fe-Fe scattering. Similarly, the Cu EXAFS analysis yields Cu-S with a coordination number of 3 and a longer-range Cu-Fe distance with a coordination number of 1, in agreement with the crystal structure (**Extended Data Table 2**). It should be noted that because of the presence of heavier elements like sulfur at about a similar distance from the metals as lighter elements like oxygen, their scattering is obscured by the Cu-S scattering and cannot be deconvoluted with EXAFS. Introduction of the substrate acetol does not induce any measurable perturbation in either the Fe or Cu XANES or EXAFS spectra (**Figure 3**), indicating that substrate binding does not measurably alter the geometric or electronic structures of the metal centers. In contrast, chemical reduction with dithionite produces observable changes in the Fe K-edge XANES spectra, including a slight decrease in pre-edge intensity and a concomitant increase in white-line intensity as indicated in the difference spectra inset in **Figure 3A**, consistent with reduction of one iron site within the cluster. These changes mirror the trends observed in the UV–vis spectra upon reduction. Importantly, the Cu K-edge XANES and EXAFS spectra remain entirely unperturbed under reducing conditions, indicating that the copper site is not involved in this redox process (**Figure 3C**). This selective response further supports the conclusion that reduction is localized at the iron centers of the novel cluster, while the copper center surprisingly remains spectroscopically and electronically unchanged.

To enhance our picture of the electronic structure of the novel cluster, we employed Mössbauer spectroscopy on ^57^Fe-enriched protein samples in their as-isolated state. Spectra were collected at 7 K in the absence of an applied magnetic field to directly assess the intrinsic electronic inequivalence of the iron sites. The closest homologs to the novel cluster, [4Fe-4S]^3+^ clusters, are most commonly described by two Mössbauer doublets assigned to two Fe^3+^ centers and two valence-delocalized “Fe^2.5+^” centers ^22,23^. The AceDH spectra can also be satisfactorily described within this conventional framework (**Figure 3E**). However, given the presence of a Cu(I) ion in the novel cluster, we also considered a model involving three distinct Mössbauer doublets, which provides a physically intuitive way to probe potential site-specific perturbations induced by the heterometallic center. In this alternative description, the relative intensities of the components are approximately 1:1:2, indicating three electronically distinguishable iron environments within the cluster (**Figure 3F**, **Extended Data Table 3**). Such differentiation likely arises from the presence of the nearby copper(I) center, which perturbs the electronic structure of the [4Fe-4S] cluster through metal–sulfur interactions and polarization of the surrounding Fe–S framework.

Finally, to connect the chemical properties of the new cofactor to the biological function of AceDH, we performed biochemical studies of the protein. (**Extended Data Figure 1, Figures 4A and F**). Importantly, we found that SolV AceDH is able to use SolV cytochrome *c_GJ_* (encoded by the *xoxGJ* gene Mfumv2_1185,^24^) as electron acceptor in a coupled assay with bovine cytochrome *c*. Given that the *xoxGJ* gene was slightly upregulated together with *gmcA* and *gmcB* upon growth on C3 compounds^5^ this cytochrome may be the physiological electron acceptor of AceDH. As the crystal structure of AceDH shows, the active site with the flavin is buried inside the protein, and whereas small molecules like Dichlorophenolindophenol (DCPIP) and PMS might be able to enter the substrate tunnel to accept the electrons generated by the oxidation of acetol from the flavin, a cytochrome such as cytochrome *c_GJ_* or any other typical physiological electron acceptor would not. It is more likely that the electrons are transferred from the flavin to the novel [Cu-4Fe-4S] cluster, which is close to the surface of the protein, and from there to a downstream redox partner; indeed, the edge-to-edge distance between the novel cluster and the flavin is only 10.9 Å, well within the “Moser-Dutton ruler” distance of 14 Å ^25,26^ enabling efficient electron transfer between the two. This poses the question why the enzyme would use a complicated cofactor as the novel [Cu-4Fe-4S] cluster for electron transfer, and not a simpler one. Since the binding motif for the novel cluster is absent in distant AceDHs, such as that from *Methylocella spp* and appears typical for verrucomicrobial methanotrophs (Supplemental Information, **Extended Data Figure 6)**, one possible reason could be that the novel cluster might offer a greater stabilization of the enzyme at the high temperatures in the geothermal habitat^27^.

**Figure 4.**
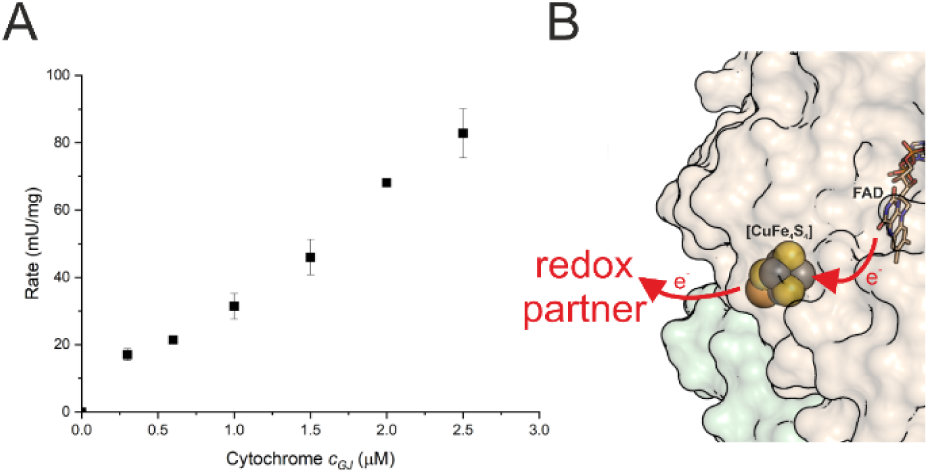
**A.** Activity assay of SolV AceDH with cytochrome *c_GJ_* and bovine cytochrome *c* as terminal electron acceptor. AceDH activity is not observed without addition of cytochrome *c_GJ_* and is linearly dependent on cytochrome *c_GJ_* concentration up to 2.5 µM. **B**. Closeup of the molecular surface of AceDH, using the color scheme from Figure 1. The novel [Cu-4Fe-4S] cluster is close to the surface and could transfer electrons from the buried flavin active site to a redox partner in the solvent.

Another possibility might be that the process requires a particular redox potential, which is offered by the novel cluster. Typically, the differentiation between ferredoxins and HiPip [4Fe-4S] cluster families is determined through solvent exposure and the protein backbone hydrogen bonding ^28–31^, but the presence of a closed-shell Cu^+^ in the [Cu-4Fe-4S] cluster could be responsible for the high potential observed in AceDH. Previously, it had been reported that addition of exogenous copper to the [3Fe-4S] cluster-containing ferredoxins of *Desulfovibrio africanus* and *Pyrococcus furiosus* results in formation of a Cu-3Fe-4S cluster, which directly influenced the reduction potential of the cluster^14,32^; however, this artificial cluster has a very different structure from the natural one presented here. EPR measurements showed this Cu to be present solely as copper(I) over the whole redox range investigated as well, like we observe here for the novel, physiological cluster in AceDH (**Figure 2**). However, selective removal of the copper through the use of various chelators was unsuccessful and we therefore could not determine the influence on the potential exactly.

In conclusion, we identified a novel heterometal FeS cluster in the form of a [Cu-4Fe-4S] cluster in the acetol dehydrogenase of *M. fumariolicum* SolV, performing an in-depth, multidisciplinary characterization, thus greatly expanding our knowledge of biology’s metallocofactor toolbox. Our findings further underscore the importance of experimental validation of protein structures and cofactors, particularly in cases where sequence-based or computational predictions may be insufficient to resolve metal site identity and composition.

## Acknowledgements

The authors wish to thank the staff of the ESRF ID23-1 beamline for their help and excellent facilities. We would also like to thank Drs. Ilme Schlichting and Mirosław Tarnawski for expert help with crystallization and for crystallographic- and XRF data collection. TRMB is very grateful to Dr. Ilme Schlichting for continuous support over many years. The authors wish to thank the I20 beamline (and staff) at Diamond Light Source for the help during XAS data collection (Proposal SP41629-1) and also Derya Demirbas for help during Mössbauer data collection. LJD thanks the SFB1309 (Deutsche Forschungsgemeinschaft (DFG) - Projektnummer 325871075, GC-MS instrument), ERC Starting Grant Lanthanophore 945846 (SolV cultivation infrastructure) and FOR BioOxCat (Deutsche Forschungsgemeinschaft (DFG) - Projektnummer 445916766, ^57^Fe source). CL was supported by the State Scholarship Fund of the China Scholarship Council (CQL: 201804910641). HOdC was supported by the European Research Council (ERC Advanced Grant project VOLCANO 669371). XY thanks Marie-Curie Fellowship for postdoctoral funding. SD and KS acknowledge the Max Planck Society for funding.

## Methods

### Growth of strain Methylacidiphilum fumariolicum SolV

Strain SolV^33^ was grown on 2-propanol or acetone with slight modifications from the procedure described by Liu et al.^5^. SolV was grown in a continuous culture with a 10 L working volume, maintained at 55 °C, stirring at 1000 rpm. The growth medium consisted of 0.2 mM MgCl_2_·6 H_2_O, 0.2 mM CaCl_2_·2 H_2_O, 1 mM Na_2_SO_4_, 2 mM K_2_SO_4_, 4 mM (NH_4_)_2_SO_4_ and 1 mM NaH_2_PO_4_·H_2_O. A solution of trace elements was added, resulting in the following final concentrations: 1 μM NiCl_2_, CoCl_2_, MoO_4_Na_2_, ZnSO_4_ and CeCl_3_; 5 μM MnCl_2_ and FeSO_4_; 10 μM CuSO_4_. The pH was set to 3 with 1 M H_2_SO_4_. 2-propanol or acetone was added to a final concentration of 13 mM and the growth medium was added at a flow of 375 ml·h^-1^ (D = 0.0375 h^-1^). The level was kept at 10 L using a level controller and the effluent was used to harvest biomass. O_2_ concentrations were kept at 20% air saturation.

To obtain ^57^Fe biomass, a trace element solution without Fe was prepared. SolV was grown in batch in bioreactor with a 10 L working volume using the same medium composition, except for iron. ^57^Fe was added in the form of a 10 µg/ml Specpure™ plasma standard solution in 0.1 M HCl (ThermoFischer) to a final concentration of 2.6 µM. 2-propanol was added in 6.5 mM steps, whenever O_2_ consumption decreased, until a final OD_600_ of 1.8-2 was achieved, after which the biomass was harvested.

SolV cells were harvested by centrifugation at 10,000 x g, 1 h 4 °C. The resulting pellet was washed once with 20 mM KPi pH 6.5 and resuspended in the same buffer in a 1:1 ratio of buffer to cell pellet. This was repeated once more and cells were snap frozen in liquid nitrogen in a 1:1 cell pellet to 20 mM KPi pH 6.5 ratio and stored at −80 °C.

### Purification

2-propanol grown SolV cells were thawed on water after which 200 mg of lysozyme was added. 20 mM KPi pH 6.5 was added to reach final volume of 100 ml and cells were homogenized 3 times using a potter homogenizer. 5 ug of DNAse I was added and cells were broken by passing them through a cell disrupter operated at 1 kbar three times. The resulting cell extract was pottered 3 times and subjected to ultracentrifugation at 150 000 g 1h 4 °C. The resulting supernatant containing the soluble proteins was loaded onto a 70 ml SP Sepharose XL column run at 5 mL/min and equilibrated in 20 mM KPi pH 6.5. After the flow through, SolV AceDH was eluted with a step of 24 mM NaCl in 20 mM KPi pH 6.5. The fraction was concentrated on a 30 kDa spin filter and buffer exchanged to 20 mM KPi 150 mM NaCl pH 6.5. This sample was then applied on a Superdex 200 10/300 GL column equilibrated with 20 mM KPi 150 mM NaCl pH 6.5 and operated at a flow of 0.5 mL/min. AceDH eluted at ∼ 15 ml. SolV AceDH was concentrated on 30 kDa spin filters and stored at −80 °C until use. All steps were performed aerobically and all chromatography steps were performed in a cold room at 7 °C.

### Gel electrophoresis and MALDI-ToF MS

SDS-PAGE^34^ was performed on AceDH by denaturing the protein sample in a SDS sample buffer solution (62.5 mM Tris-HCl pH 6.8, 2% SDS, 10% glycerol, 5% β-mercaptoethanol, 0.005 % bromophenol blue) at 100 °C for 10 min. The sample was then run on a 4-16% SDS gel at 100 V until the dye front reached the end of the gel. A PageRuler^TM^ Plus Prestained Protein Ladder (Thermo Fisher Scientific) was used as a MW indicator. Gels were stained with Coomassie Brilliant blue. Bands were cut out from the SDS PAGE gel and digested with trypsin for protein identification using MALDI-ToF MS analysis according to a protocol by Farhoud et al.^35^ Spectra were collected using a Microflex LRF MALDI-TOF (Bruker Daltonic) and analyzed using the Mascot Peptide mass Fingerprint program against an in-house *Methylacidiphilum fumariolicum* SolV protein database, with a peptide tolerance of 0.3 Da, allowance of one missed cleavage and methionine oxidation as a variable modification

### ICP-MS

Metal content was analysed by ICP-MS on a Series I ICP MS (Thermo Scientific, Breda, the Netherlands). For detecting metals, 450 μg of purified protein was washed with ddH_2_O (total volume used, 1 mL) on a Vivaspin™ 500 Centrifugal Concentrator (Sartorius, Germany) and destructed with nitric acid (10%) at 90°C for 30 to 60 min and diluted to 5 ml with water (final nitric acid concentration 1%).

### Activity assay

Acetol dehydrogenase activity was measured spectrophotometrically following the reduction of 2,6-dichlorophenolindophenol (DCPIP) as redox dye at 600 nm, coupled with phenazine ethosulfate (PES) as electron shuttle^36^. The assay was performed at 45 °C in a 1 cm path length Suprasil quartz cuvette (Hellma) using a Cary 60 spectrophotometer (Agilent). Each reaction contained a total volume of 200 µl, including 10 mM phosphate buffer (pH 7.4), 2 mM PES, and 1 mM DCPIP. To minimize background reactions, an assay premix containing 4 mM PES and 2 mM DCPIP in buffer was prepared, heated to 45°C for 15 min, and then stored in an amber Falcon tube on ice to prevent light-induced degradation. Acetol concentrations were varied between 1-50 µM and after establishing a background, reactions were started by the addition of 3.2 µg SolV AceDH. Initial slopes were determined and fitted to the Michaelis Menten equation to obtain enzyme kinetic parameters. To determine the pH optimum the assay was performed in a multicomponent buffer consisting of 2.5 mM citric acid, 2.5 mM Bis-Tris, 2.5 mM Tris and 2.5 mM CHES in a range of 5.2-9 at 45 °C. To determine the temperature optimum, the assay was performed in the same multicomponent buffer at temperatures in the range of 25-95 °C. In both cases an acetol concentration of 500 µM was used and DCPIP extinction coefficients in the various buffer systems were taken from Jahn et al.^36^

To examine the role of cytochrome *c_GJ_* as electron acceptor, acetol dehydrogenase activity was determined with an Epoch 2 plate reader by following the reduction of bovine cytochrome *c* at 550 nm. The assay was performed at 45 °C in a 96 well plate. Each reaction contained 200 µl total volume, including 50 mM Pipes pH 7, 50 µM cytochrome *c*, 50 µM acetol, 100 nM SolV AceDH and 0-2.5 µM cytochrome *c_GJ_*. Reactions were performed in triplicate and the absorbance was pathlength corrected using 977/900 nm absorbance. After recording the initial absorbance, reactions were started by addition of acetol. Initial slopes were determined and the rates were plotted versus the cytochrome *c_GJ_* concentration.

#### Product identification

To detect the reaction product methylglyoxal, gas chromatography coupled with mass spectrometry (GC-MS) was used. For this, a derivatization protocol for methylglyoxal was adapted from the literature^37,38^. All reactions were run in 10 mM potassium phosphate (KP) buffer pH 7.4 (CAS 7778-53-2, Sigma-Aldrich, reagent grade, 98%) and the total reaction volume was 500 µL. 100 mM PES (10510-77-7, ≥95%, Sigma-Aldrich, in MilliQ) was used as stock. 1 M acetol (116-09-6, 90%, Sigma-Aldrich) solution in KP buffer was used as stock. A 40% methylglyoxal solution in water was used as stock. The derivatization reagent O-(2,3,4,5,6-pentafluorobenzyl)hydroxylamine hydrochloride (PFBOA, 57981-02-9, ≥98%, Sigma-Aldrich) was used as solid and directly added to the samples. An AceDH batch that was heated to 60°C in air for 45 min was used. All samples and controls were heated in 2 mL Eppendorf vials to 50°C for one hour in a heating block (PFBOA not yet added). Subsequently samples were centrifuged for 1 min to settle any precipitates and the liquids transferred to 15 mL Falcon tubes (VWR). 4 mL of a 1 mM HCL solution was added and 5 mg PFBOA (Sigma-Aldrich) added (if used). The samples were vortexed until the white solid of PFBOA was completely dissolved and subsequently incubated at 50°C for 30 min in a heating block. Once cooled to 25°C, approximately 0.5 g NaCl (ACS Reag. Ph. Eur. VWR Chemicals) was added to all samples and vortexed until the NaCl had dissolved. 3 mL of a 1:1 mixture of diethylether and n-hexane were added to the Falcon tubes and the samples briefly vortexed to extract the analytes. Approximately 2.75 mL of the organic layer were transferred into a second 15 mL Falcon tube and concentrated to approximately 500 µL using a stream of nitrogen. The samples were filtered over 0.2 μm PTFE filters and subjected to GC-MS analysis. The samples were injected (1 µl, split 100:1) onto an Agilent® 7920 GC equipped with a 30 m HP5-MS column (Agilent® 19091S433UI) coupled to an Agilent® 5970 EI mass spectrometer. The injector temperature was set to 280 °C and the temperature of the ion source to 230 °C. The initial oven temperature was 80 °C, then held there for 2 min, then ramped to 150 °C at 15 °C/min, and finally ramped to 280 °C at 30°C/min and held there for 8 min. Mass spectra were recorded in scan mode between 70-400 m/z. Methylglyoxal bis-(O-pentafluorophenylmethyl-oxime) eluted between 10.4-10.7 min and was identified by comparison to the NIST database. Without derivatization, methylglyoxal could not be detected. It was also ruled out that impurities in acetol (90%, technical grade) contained detectable levels of methylglyoxal.

### UV-Vis Absorption spectroscopy

UV-Vis spectra were recorded in a range of 250-700 nm with an Agilent Cary60 spectrophotometer using 1 cm Suprasil Quartz cuvettes (Hellma). Spectra were recorded aerobically at room temperature or in an anaerobic hood maintained at 17 °C. After obtaining as isolated spectra, samples were reduced by addition of solid sodium dithionite powder, 1 equivalent of acetol or 1 equivalent of sodium ascorbate in the presence of 200 nM phenazine methosulfate (PMS). SolV AceDH concentrations were determined using the absorbance at 280 nm and an extinction coefficient of 108,750 M^-1^·cm^-1^ determined from the GmcA & GmcB amino acid sequence using the Expasy prot param webserver.

### Temperature-dependent Circular Dichroism spectroscopy

CD spectra were collected on a JASCO J-1500 spectropolarimeter (JASCO Corporation, Tokyo, Japan) equipped with a Peltier-type temperature-control unit. All measurements were performed in the far-UV region, scanning from 190-260 nm with a data pitch of 0.1 nm. The scan speed was set to 100 nm min^−1^. Measurements were performed using a 1 mm path-length quartz suprasil cuvette (Hellma). Three detection channels were recorded simultaneously: ellipticity θ (mdeg), high-tension (HT) voltage (V), and UV absorbance. Spectral regions where the HT voltage exceeded 700 V were excluded from analysis, typically occurring at wavelengths below ∼ 200 nm in the present buffer system. A buffer blank (20 mM KP, 150 mM NaCl, pH 7) without protein was used for temperature-resolved baseline correction and a 2 μM AceDH protein concentration was used. Full-wavelength spectra (190-260 nm) were recorded at 13 discrete temperatures during a heating ramp: 15, 20, 25, 30, 35, 40, 45, 50, 55, 60, 65, 70, and 75°C (5°C increments). The sample temperature was increased at a rate of 5°C min^−1^, with an equilibration period of 20 s at each target temperature prior to spectrum acquisition. Following the heating phase, the sample was cooled from 75 back to 15°C using the identical 13-point step protocol (cooling ramp), to assess the reversibility of any thermally induced structural changes. Buffer blank spectra were collected under the identical thermal program and baseline-corrected spectra were obtained by subtracting the temperature-matched buffer spectrum from the corresponding sample spectrum at each wavelength.

### EPR spectroscopy

EPR spectroscopy was performed on an Elexsys E500-II, equipped with an X-band (∼9.5 GHz) and Q-band (34 GHz) FT-EPR microwave bridges. For X-band measurements, samples were measured at low temperatures using an ESR 900 cryostat (Oxford Instruments) and a Mercury ITC nano temperature controller (Oxford Instruments), paired with either a Bruker dual-mode TE102 cavity or a Bruker SHQE resonator. Samples were prepared in custom quartz tubes (3.8 mm OD, 0.5 mm wall thickness) or, for the electrochemical titration, 15 μL (200 µM AceDH) aliquots were placed in 2 mm OD x ∼3 cm (length) quartz EPR tubes (Wilmad) and placed in a larger EPR tube for measurement. Approximately 100 μL of water was added to the bottom of the tube, surrounding the inner quartz EPR tube, to improve heat transfer of the sample with its surroundings. All samples were frozen in liquid nitrogen and stored in liquid nitrogen until measurement. Q-band samples were prepared in custom 1.6 mm (OD) quartz tubes and measured in a Q-band resonator (ER 5106QT Bruker) within an Oxford CF935 cryostat controlled by the same temperature controller above. Parameters of the collected EPR spectra are provided in the associated figure captions. All EPR data was processed and analyzed in Matlab 2025a. EPR spectra were reproduced by simulation via the open-source EasySpin (v. 6.0.11) package for Matlab. All spin Hamiltonian parameters are provided in the main text.

### Redox titration

Optical redox titrations were performed aerobically using a custom made optically transparent thin layer electrochemical cell^39^ with Suprasil Quartz glass windows and a pathlength less than 10 µm. Goldgrid working electrodes were modified by boiling for 10 min in a 1 mg/ml PATS-3 solution. A platinum wire was used as a counter electrode and a Ag/AgCl wire in 3 M KCl was used as a reference electrode, calibrated for SHE by titration of bovine heart cytochrome *c.* The cell was connected to a Metrohm autolab PGSTAT204 potentiostat and spectra were recorded using a Cary 60 spectrophotometer in the range of 250 to 700 nm. Titrations were performed on 1.4 mM AceDH in either 100 mM Tris pH 8 or 100 mM Tris pH 7, supplemented with 150 mM KCl and 40 µM mediator mixture (ferricyanide (E^0^′ = +430 mV); 1,4-benzoquinone (E^0^′ = +280 mV); 2,5-dimethyl-1,4-benzoquinone (E^0^′ = +180 mV); 1,2-naphthoquinone (E^0^′ = +145 mV); phenazine methosulfate (E^0^′ = +80 mV); 1,4-naphthoquinone (E^0^′ = +60 mV); phenazine ethosulfate (E^0^′ = +55 mV); 5-hydroxy-1,4-naphthoquinone (E^0^′ = +30 mV); 1,2-dimethyl-1,4-naphthoquinone (E^0^′ = 0 mV); 2,5-dihydroxy-p-benzoquinone (E^0^′ = −60 mV); 5,8-dihydroxy-1,4-naphthoquinone (E^0^′ = −145 mV), 9,10-anthraquinone (E^0^′ = −184 mV), 9,10-anthraquinone-2-sulfonate (E^0^′ = −225 mV); benzyl viologen (E^0^′ = −350 mV); and methyl viologen (E^0^′ = −440 mV/−772 mV).

Titrations were performed in the range of +700 to −100 mV vs SHE in 50 mV steps, with a 25 mV offset between reductive and oxidative directions. At each potential the sample was equilibrated for 10 min before a spectrum was recorded. The absorbance at 460 nm was plotted as a function of the applied potential and fitted to a sum of Nernst curves using Origin 2021b. Additionally, the spectra were analysed using the mfit-nernst function of the QSOAS software^40^ to obtain a global fit and extract the redox potentials and corresponding oxidized minus reduced difference spectra of the cofactors.

The electrochemical EPR titration was performed as a two-electrode setup in a low-volume cell kit (Prosense) equipped with a Pt counter electrode and an Ag/AgCl reference electrode with nitrogen gas purging through the headspace. The voltage potential of the cell was measured with a VersaSTAT-4 instrument (Ametek GmbH). A 1,200 µL solution of 200 μM AceDH, 100 mM Tris pH7, 150 mM KCl, and 40 µM mediators (see above) was prepared for study. The titration was performed through small additions (1 to 4 μL) of 2, 5, and 50 mM sodium dithionite or ferricyanide solutions and 15 μL aliquots were taken and prepared as described above.

### Crystal structure determination

The SolV AceDH protein was concentrated to 30mg/ml in 10 mM potassium phosphate (KPi) buffer, pH 7.4. Crystallization conditions were screened in 96-well sitting drop JCSG+ screen plates^41^ using a Mosquito pipetting robot (SPT Labtech, Jena, Germany). Initial conditions were optimized in 24-well Linbro plates using hanging drop setups. The best crystals were obtained using a reservoir solution consisting of 50 mM citric acid, 2 M LiCl and a 1 µL drop of reservoir solution was mixed with 1 µL of protein solution. The drops were equilibrated against 900 µL reservoir solution at 293 K. The crystals grew within 2 days. Native data were collected to 1.8 Å resolution at 100 K, from crystals cryoprotected by a brief soak in mother liquor supplemented with 30% ethylene glycol and few crystals of sodium dithionite, at the ID23-1 beamline of the European Synchrotron Radiation Facility (ESRF, Grenoble, France). Data were processed using XDS^42^ and were phased by molecular replacement with PHASER^43^ using an AlphaFold2 model^44^ obtained using ColabFold^10^ as the search model. The final model was obtained by iterative cycles of rebuilding using COOT^45^ and refinement using PHENIX. Data collection and model statistics are reported in Extended Data Table S1 and the structure was deposited in the Protein Data Bank^46^ under entry code 9TTU. To identify an additional atom in the iron-sulfur cluster, an X-ray fluorescence (XRF) spectrum was collected at the ID23-1 beamline of the ESRF (Grenoble, France) from a crystal kept at 100 K. Single-wavelength anomalous dispersion data were collected at the same beamline at wavelengths just above the iron- and copper K absorption edges; statistics are given in Extended Data Table S1.

### Sequence analysis

The SolV AceDH large (GmcA); mfumv2_1610) and small (GmcB; mfumv2_1609) subunits were used as query in a BlastP search, to find homologous proteins. Closely related proteins were selected and a phylogenetic analysis was performed using MEGA12^47^. To infer the evolutionary history the Neighbor-Joining method was used and the confidence of the resulting phylogenetic tree was assessed by bootstrapping analysis (500 replicates). Multiple sequence alignments were performed using the Clustal Omega webservice of the EMBL-EBI (https://www.ebi.ac.uk/jdispatcher/msa/clustalo)

### X-ray absorption Spectroscopy

#### Data collection

Fe and Cu K-edge X-ray absorption spectroscopy (XAS) measurements were performed at the I20-scanning beamline of Diamond Light Source (Oxfordshire, UK). Incident energy selection was achieved using a four-bounce Si(111) monochromator, and rhodium-coated mirrors were employed for harmonic rejection, providing an unattenuated photon flux of approximately 1 × 10^12^ photons s^-1^ at the sample position. The X-ray beam was focused to a spot size of ∼0.3 × 0.4 mm² (vertical × horizontal, full width at half maximum). The NifH protein, which is the reductase protein component of Mo-Nitrogenase, was purified from *Azotobacter vinelandii* following a previously established purification protocol^48^. In order to make sure the sample was in the resting +1 oxidation state, 2.5 mM dithionite was added to the purified sample. XAS of a 1 mM protein sample was measured similarly as described below for the AceDH samples.

AceDH protein samples (∼ 0.75-1 mM protein concentration), under different conditions (prepared similarly as described before), were mounted in a top-loading exchange-gas pulse-tube helium cryostat and maintained at 10 K throughout all measurements to minimize beam-induced damage. Beam damage was assessed prior to data collection using a series of short, low-resolution near-edge scans under progressively attenuated flux. Based on these tests, the total flux of the incident beam was attenuated using 2.5 mm C film for both Fe and Cu K-edge measurements to ensure spectral stability during each ∼15 min scan. Full spectra were collected from fresh sample positions for every scan.

For the Fe K-edge, spectra were acquired over 6909-7859 eV and calibrated by simultaneous measurement of an Fe foil reference, with the first inflection point of the Fe foil set to 7111.2 eV. For the Cu K-edge, incident energies were scanned from 8859 to 9840 eV, with calibration performed using a Cu foil for which the first inflection point was set to 8980.3 eV. Three ionization chambers were positioned along the beam path to record incident (I_0_), transmitted (I_t_), and reference (I_ref_) intensities. Fluorescence signals from the sample were collected using a 64-element monolithic Ge detector, with readout processed by an Xspress4 digital pulse processor.

#### Data analysis

X-ray absorption spectroscopy (XAS) data were processed using the Demeter software package^49^. Data normalization was carried out by applying a linear regression to the pre-edge region and a cubic polynomial regression to the post-edge region. Background subtraction was performed using a spline function over the post-edge region, with a *k*-range of 0-14 Å^-1^, an *R*-background value of 1.0 Å, and a *k*-weight of 3 for both Fe and Cu. The extended X-ray absorption fine structure (EXAFS) signal was extracted and Fourier transformed using a Hanning window over the *k*-ranges of 2-13.5 Å^-1^ for Fe and 2-11 Å^-1^ for Cu. Modeling and fitting of the Fourier-transformed EXAFS (FT-EXAFS) data were performed using the Artemis module with embedded FEFF6 calculations. Scattering paths were generated based on the resolved crystalline structure of AceDH. The atomic clusters used for the FEFF calculations are shown in Extended Data Figure 4. Scattering paths with similar effective path lengths (ΔR ≤ 0.14 Å for Fe and ≤ 0.17 Å for Cu) were grouped and treated as degenerate during fitting. For the Cu FT-EXAFS analysis, it is noted that backscattering from a single O atom may be partially masked by multiple single-scattering contributions from neighboring S atoms. Therefore, no Cu-O path is included in the fitting. Fitting was conducted over a non-phase shift corrected *R*-range of 1.0-3.0 Å for Fe and 1.2-2.8 Å for Cu. The amplitude reduction factor (S_0_^2^) was fixed at 0.9 for all scattering paths. Initial fits were performed with fixed path degeneracies and path length derived from the optimized local structure, allowing the Debye-Waller factor (σ^2^) and energy shift (E_0_) to vary. In the final refinement, all fitting parameters, including path length, σ^2^, and E_0_, were allowed to float freely. The final fitting results are summarized in **Extended Data Table 2**, and representative fits are shown in Extended Data Figure 4.

#### Mössbauer Spectroscopy

^57^Fe Mössbauer spectra were recorded at 7K with a spectrometer using a Janis Research (Wilmington, MA) SuperVaritemp dewar. Isomer shifts are quoted relative to the α-Fe metal at 298 K. Mössbauer spectral simulations were performed using the local program Mfit (*mf*, written by Dr. Eckard Bill) using the minimum number of necessary quadrupole doublets to gauge the average isomer shift of each spectrum.

## Supplementary information and Extended Data Figures and Tables

### Properties of AceDH

Through an activity-based purification process, AceDH from *M. fumariolicum* SolV was isolated from the soluble fraction of a cell lysate using two consecutives chromatography steps. SDS_PAGE analysis (**Extended Data Figure 1A**) revealed two prominent protein bands, which were identified by MALDI-TOF MS as the products of gmcA (Mfumv2_1610) and gmcB (Mfumv2_1609). The gmcA gene encodes a 526 amino acid flavin adenine dinucleotide (FAD)-dependent FAD oxidoreductase belonging to the glucose-methanol-choline (GMC) enzyme family. Domain analysis using Pfam identified conserved GMC oxidoreductase N-terminal (GMC_oxred_N; PF00732, residues 123 to 298) and C-terminal (GMC_oxred_C; PF05199, residues 397 to 512) domains (Supplementary see alignments below). The two genes are part of a 13-gene cluster that was highly upregulated during growth on C3 compounds compared to C1 compounds in both *M. fumariolicum* SolV4 and *M. caldifontis* IT623 and postulated to be involved in acetol oxidation to methylglyoxal, an activity that was previously confirmed in *M. sylvestris* BL224. In kinetic experiments, AceDH was found to oxidize acetol with a *k_cat_* of 0.69 ± 0.01 s^-1^, and a *K_M_* of 2.3 ± 0.2 µM in an assay with DCPIP as the terminal electron acceptor and PMS as electron shuttle (**Extended Data Figure 1B**). Production of methylglyoxal was confirmed through GC-MS analysis (**Extended Data Figure 5, Extended Data Table 4**). The enzyme’s pH optimum was determined to be at 8.1 **(Extended Data Figure 1C**) and its optimal temperature was 80 °C. (**Extended Data Figure 1C**) AceDH, being isolated from a thermophilic microorganism, retained ∼50% of its maximum activity at 95 °C and in CD spectrometry showed no major unfolding between 15 and 75°C (**Extended Data Figure 1F**). Higher temperatures could not be investigated with circular dichroisms spectroscopy as the concentrations necessary for CD measurements (2 μM) lead to partial precipitation of AceDH above 80 °C. Production of methylglyoxal was confirmed through GC-MS analysis after derivatization of the reaction mixture with O-(2,3,4,5,6-penta-fluorobenzyl)hydroxylamine hydrochloride (PFBOA). SolV AceDH did not show any activity on other tested alcohols (methanol, 1-propanol, 2-propanol, and 1,2-propanediol).

The electron density maps of the AceDH crystal structure are of excellent quality. They indicate that in the crystal structure, the FAD CBM atom is covalently bonded to the His91 NE2 atom. (**Extended Data Figure 1G**). The apparent active site further contains two crystallographic water molecules close to the flavin, hydrogen-bonded to Thr451α, His458α, and Asn501α on one side of the active site cavity. His458α is hydrogen bonded to His297α, which displays unfavorable Ramachandran angles, though its density is perfectly defined (**Extended Data Figure 1H**). The side chain of His297α is hydrogen bonded to three backbone CO groups. Two of these, from Gly497α and Ala 498α interact with the NE2 atom while the third, from Glu500α, is at only 3.1 Å distance from the CE1 atom, suggesting a CH···O hydrogen bond. The other side of the active site is predominantly hydrophobic, with Met318α and Phe194α forming the wall of the cavity.

### Catalytic mechanism

Attempts to cocrystallize AceDH with acetol or to soak acetol into the crystals were unsuccessful. Nevertheless, the structure does allow some features of a catalytic mechanism to be proposed. The oxidation of acetol to methylglyoxal requires the abstraction of two protons and two electrons from the substrate. In AceDH, the electrons would be transferred to the flavin, but at least one of the protons would need to be accepted by another moiety. If the carbon atom displaying the hydroxyl group of acetol is positioned close to the flavin N5 atom (which would put the substrate’s methyl group on the hydrophobic side of the cavity), the N5 could abstract one of its protons. In that situation, the hydroxyl group would be close to the His458α NE2 atom, which could abstract its proton. As mentioned above, His458α is hydrogen bonded through its ND1 to His297α, which is involved in an unusual CH···O hydrogen bond. This could function as a charge-relay system, as was suggested for the histidine in Ser-His-Asp proteases and -esterases ^50^, removing the positive charge of the abstracted proton from the active site and storing it on His297α, possibly to aid the abstraction of a second electron and/or proton (**Extended Data Figure 1I**).

### Details of the EPR spectra

Inspection of the coordination sphere of the [Cu-4Fe-4S] cluster (**Figure 1**) does not reveal any clear origins of the hyperfine splitting (e.g. coordinated ^14^N, *I* = 1, of histidine or arginine residues). Given the narrow EPR linewidth of the *g*_∥_ feature, it is plausible a weak hyperfine coupling of the copper ion and its two isotopes (^63^Cu 69.17%, ^65^Cu 30.83%, both *I* = 3/2) could be resolved. Simulation of the X-band EPR spectrum with an axial g-tensor and the inclusion of a single 23 MHz copper hyperfine coupling interaction along *g*_∥_ reproduces the general shape of the second derivative line shape; however, it fails to reproduce the complete multiline pattern, suggesting more than one nucleus is contributing. Simulation with a 20 MHz proton (^1^H) coupling further splits the copper hyperfine pattern and reproduces it more closely. Such a large ^1^H hyperfine coupling is considered a bit unusual as a 22 MHz proton is more typical of a coordinated hydride^51,52^ than the more probable candidate of β-cysteinyl protons, which may have couplings upwards of 10 MHz as demonstrated in [4Fe-4S] model complexes^15,53^. However, such a 22 MHz ^1^H is more suggestive of large and/or direct proton spin polarization, such as the protonation of a µ_3_-sufide or the formation of FeS-cysteinyl radical structure, but hyperfine measurements in such examples are rare thus far. Similarly, a ^14^N coupling interaction of ∼15 MHz, also splits the edges of the *g*_∥_ feature appropriately, improving the fit. As alluded to earlier, the crystal structure suggests no possible origin of a strongly coupled ^14^N nucleus. The final evaluated possibility is that the feature is the result of the superposition of two spectra, each with similar copper hyperfine couplings but the *g*_∥_ values differ slightly, sufficient to produce the complex hyperfine pattern. The simulation of this scenario reproduces the X-band spectrum well; however, as discussed below, the higher-frequency Q-band data eliminates this possibility (**Extended Data Figure 3**).

The higher-frequency Q-band (34 GHz) EPR spectrum of AceDH also exhibits an axial spectrum, with a fairly sharp *g*_∥_ feature and a broader *g*_⊥_. The *g*_∥_ feature lacks resolved hyperfine features, and the second derivative of this position does not show the highly resolved features observed in X-band (**Extended Data Figure 3**). The increased EPR line broadening at microwave-frequency (and magnetic field strength) suggests possible g-strain (distribution of the g-value), as often observed for FeS clusters, explaining the loss of resolved hyperfine features.

While the simulation of the Q-band EPR spectrum by a single species, regardless of hyperfine contributions, reproduces the spectrum well, the data analysis clearly eliminates the possibility of two components. As evaluated for the X-band EPR (**Extended Data Figure S3**), the simulation of two spectral components with similar Cu hyperfine couplings but different *g*_∥_ values is clearly unsatisfactory for accurately reproducing the Q-band experimental data (**Extended Data Figure S3**). Therefore, the X-band analysis scenarios of [Cu,^14^N] or [Cu,^1^H] reproduce the data best.

**Extended Data Figure 1.**
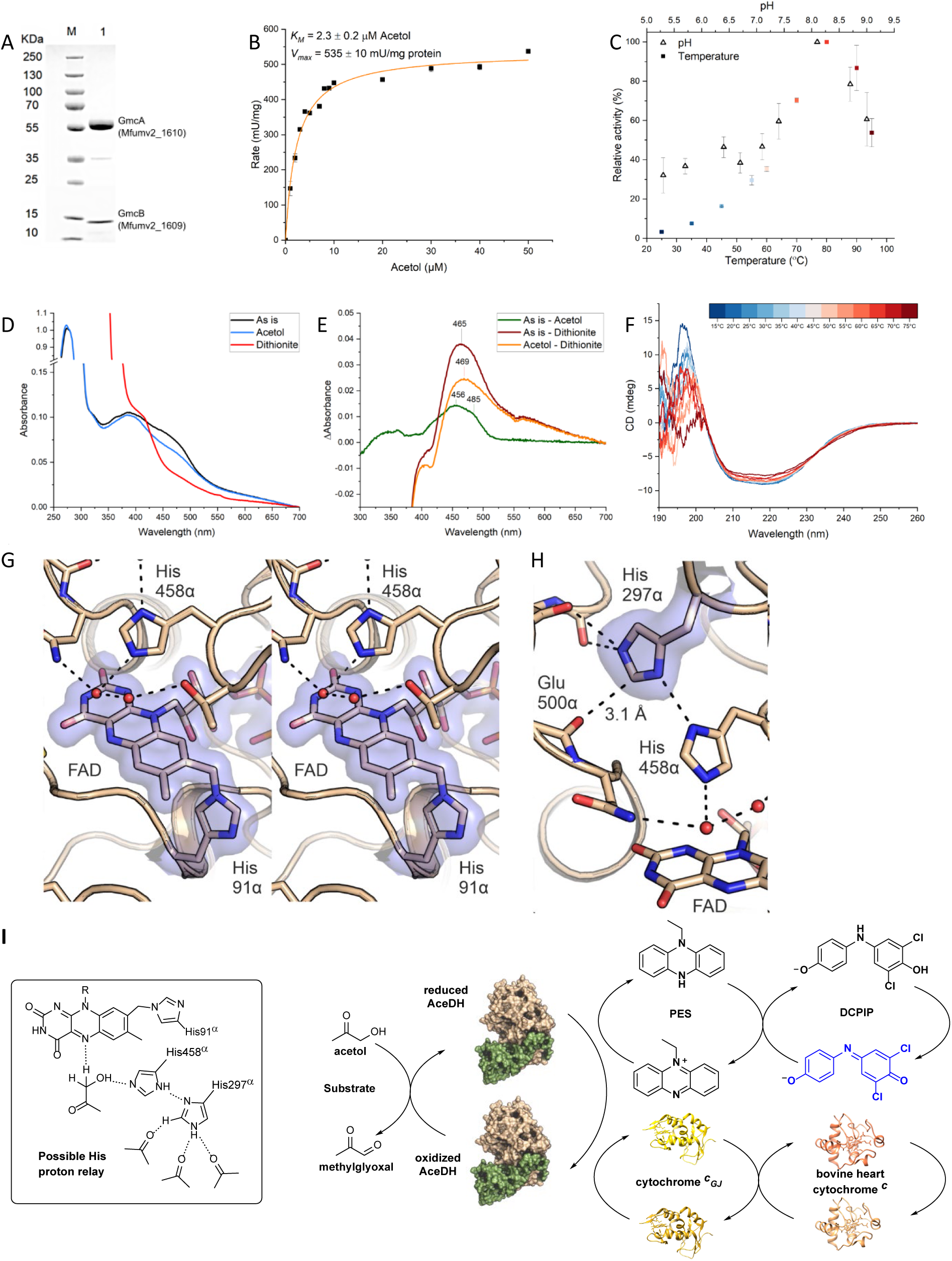
Properties of AceDH. **A.** SDS Page of acetol dehydrogenase isolated from *M. fumariolicum* SolV. The bands at 58 kDa and 14 kDa were identified through MALDI-ToF MS analysis of their tryptic digests as gene products of *gmcA (Mfumv2_1610)* and *gmcB (Mfumv2_1609)*, respectively. **B.** Michaelis-Menten kinetics of AceDH oxidizing acetol. **C.** Temperature- and pH-dependence of AceDH activity. **D.** UV-Vis spectra of AceDH in the as-isolated state and after treatment with acetol and sodium dithionite. **E.** Difference spectra calculated from the data shown in panel D. **F.** Circular dichroism spectra of AceDH at various temperatures. **G.** Stereofigure showing the active site of AceDH with the protein shown as a cartoon and the FAD and selected residues shown as sticks. The final, refined 2*m*F_o_-DF_c_ electron density map (contoured at 1 σ) is shown as a transparent blue surface for the FAD and the covalently linked αHis91. **H.** Another closeup of the catalytic site, centered on αHis297 which forms an unusual CH…O hydrogen bond to the αGlu500 carbonyl oxygen. **I.** Possible His-Proton-relay pathway and the two described assay setups of AceDH.

**Extended Data Figure 2.**
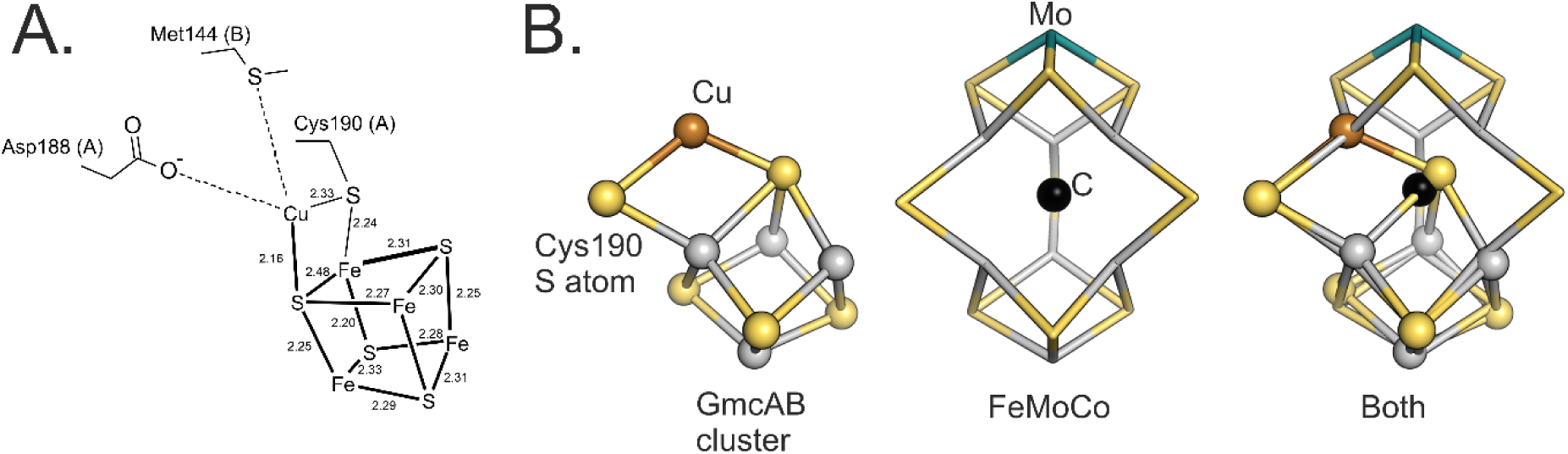
**A.** Detailed structure of the novel cluster from the A molecule of the crystal structure. Interatomic distances are shown in Ångstroms. **B.** Comparison of the atomic positions in the novel cofactor with those in the nitrogenase FeMo cofactor. The novel cofactor (left panel) and the FeMoCo (middle panel) can be superimposed (right panel) to illustrate that the distances and angles in the novel cofactor are chemically plausible for a metallocofactor with multiple metal- and sulfur atoms.

**Extended Data Figure 3.**
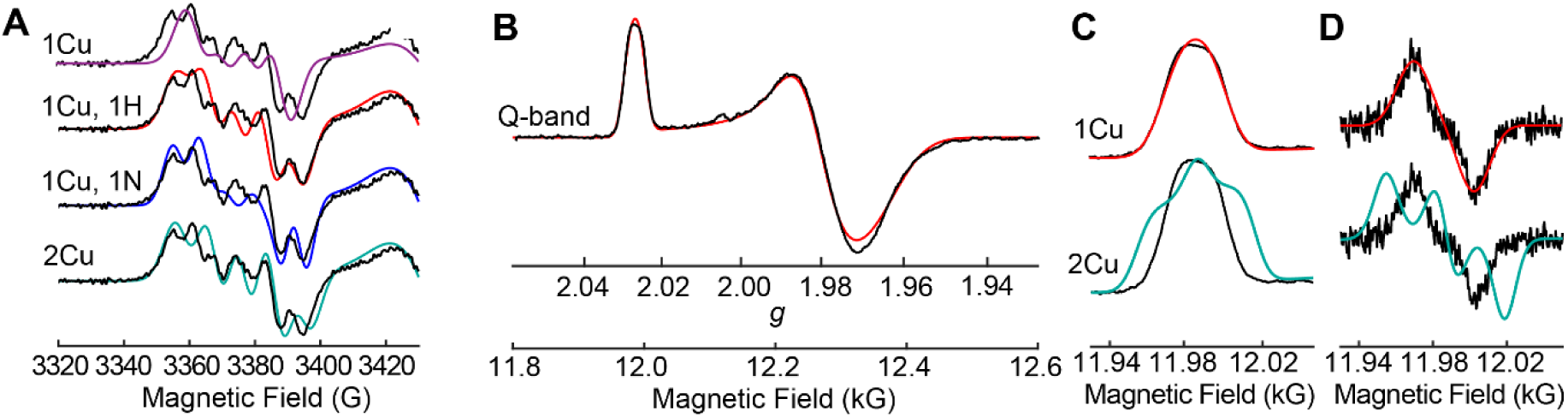
**A.** Second derivative X-band EPR signal of the g_∥_ region (black) with associated simulations of different nuclei pairs. For “1Cu,” “1Cu,1H,” and “1Cu,1N” a constant A(Cu) = 22.7 MHz was used and ^1^H or ^14^N coupling adjusted. The “2Cu” has two species with slightly different *g*_∥_ values and similar copper hyperfine couplings. **B.** Q-band EPR of SolV AceDH (black) and simulation (red), including only a single Cu nucleus. **C.** Expansion of the g_∥_ region and **D.** the first derivative of the signal, with simulations. A constant A(Cu) = 22.7 MHz was used. The “2Cu” has two species with slightly different g_∥_ values and similar copper hyperfine couplings.

**Extended Data Figure 4.**
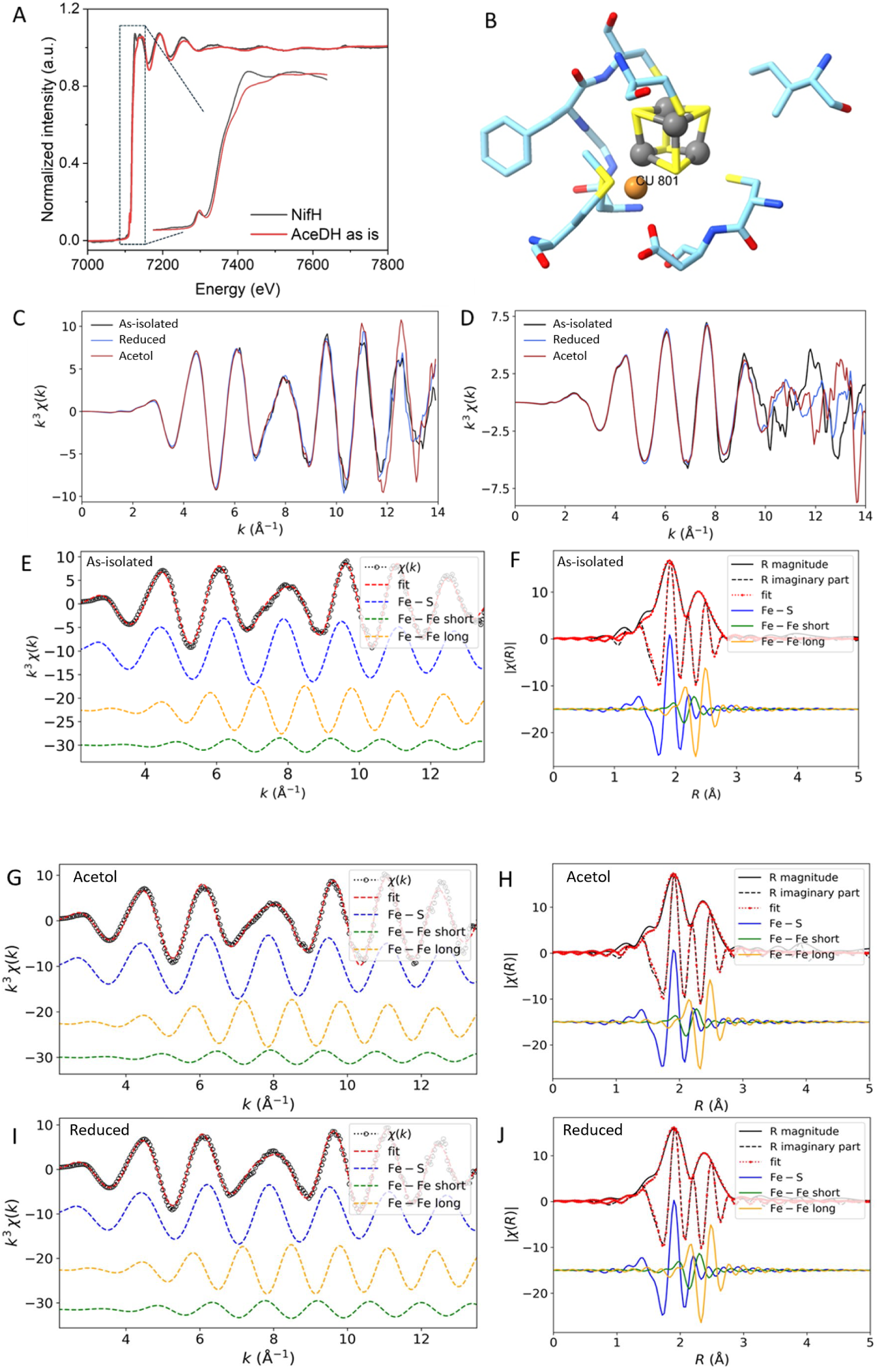
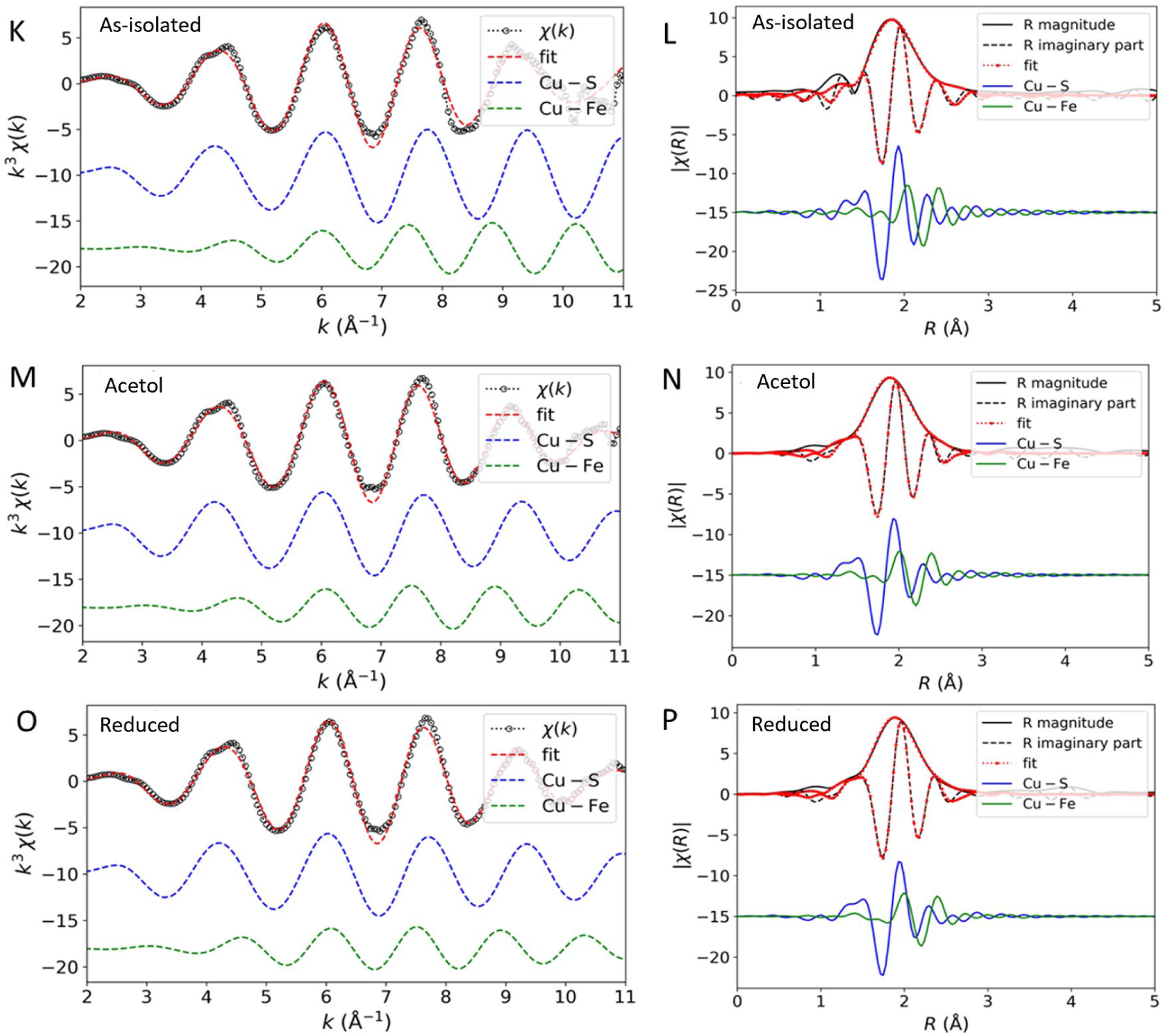
**A.** Fe K-edge XAS of NifH and AceDH in the as-isolated resting state, with the inset showing zoomed-in XANES region of the spectra. **B**. Local structure used for Feff calculations (obtained from the crystal structure). **C**. *k*^3^-weighted k-space EXAFS spectra of all protein samples at the Fe K-edge and **D.** the Cu K-edge. **E.** Fe K-edge EXAFS fitting results for as-isolated AceDH, showing the total fit and individual path contributions in *k*^3^-weighted χ(k) and its Fourier-transformed spectra (magnitude and imaginary components) (**F**). **G**. Fe K-edge EXAFS fitting results for acetol adsorbed AceDH, showing the total fit and individual path contributions in *k*^3^-weighted χ(k) and its Fourier-transformed spectra (magnitude and imaginary components) (**H**). **I.** Fe K-edge EXAFS fitting results for reduced AceDH, showing the total fit and individual path contributions in k^3^-weighted χ(k) and its Fourier-transformed spectra (magnitude and imaginary components) (**J**). **K.** Cu K-edge EXAFS fitting results for as-isolated AceDH, showing the total fit and individual path contributions in *k*^3^-weighted χ(k) and its Fourier-transformed spectra (magnitude and imaginary components) (**L**). **M.** Cu K-edge EXAFS fitting results for acetol adsorbed AceDH, showing the total fit and individual path contributions in *k*^3^-weighted χ(k) and its Fourier-transformed spectra (magnitude and imaginary components) (**N**). **O.** Cu K-edge EXAFS fitting results for reduced AceDH, showing the total fit and individual path contributions in *k*^3^-weighted χ(k) and its Fourier-transformed spectra (magnitude and imaginary components) (**P**).

**Extended Data Figure 5.**
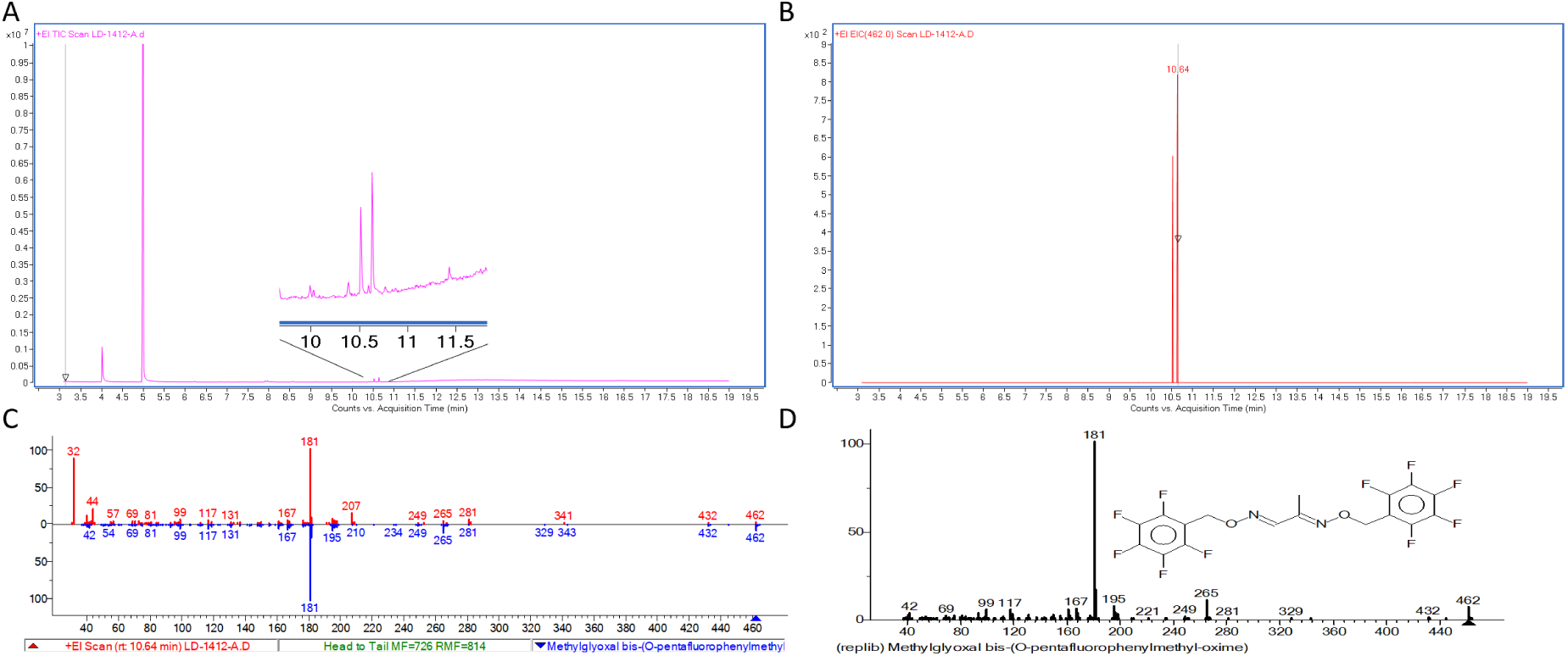
Detection of methylglyoxal as product. **A** GC-MS trace of LD-1412-A sample with inset the region of methylglyoxal bis-(O-pentafluorophenylmethyl-oxime). **B** Extracted ion chromatogram for the mass of methylglyoxal bis-(O-pentafluorophenylmethyl-oxime), m/z 462. **C** Comparison of fragmentation pattern at rt 10.54 min (top, red) with NIST database data (bottom, blue). **D** Fragmentation pattern for target molecule from NIST database

**Extended Data Figure 6.**
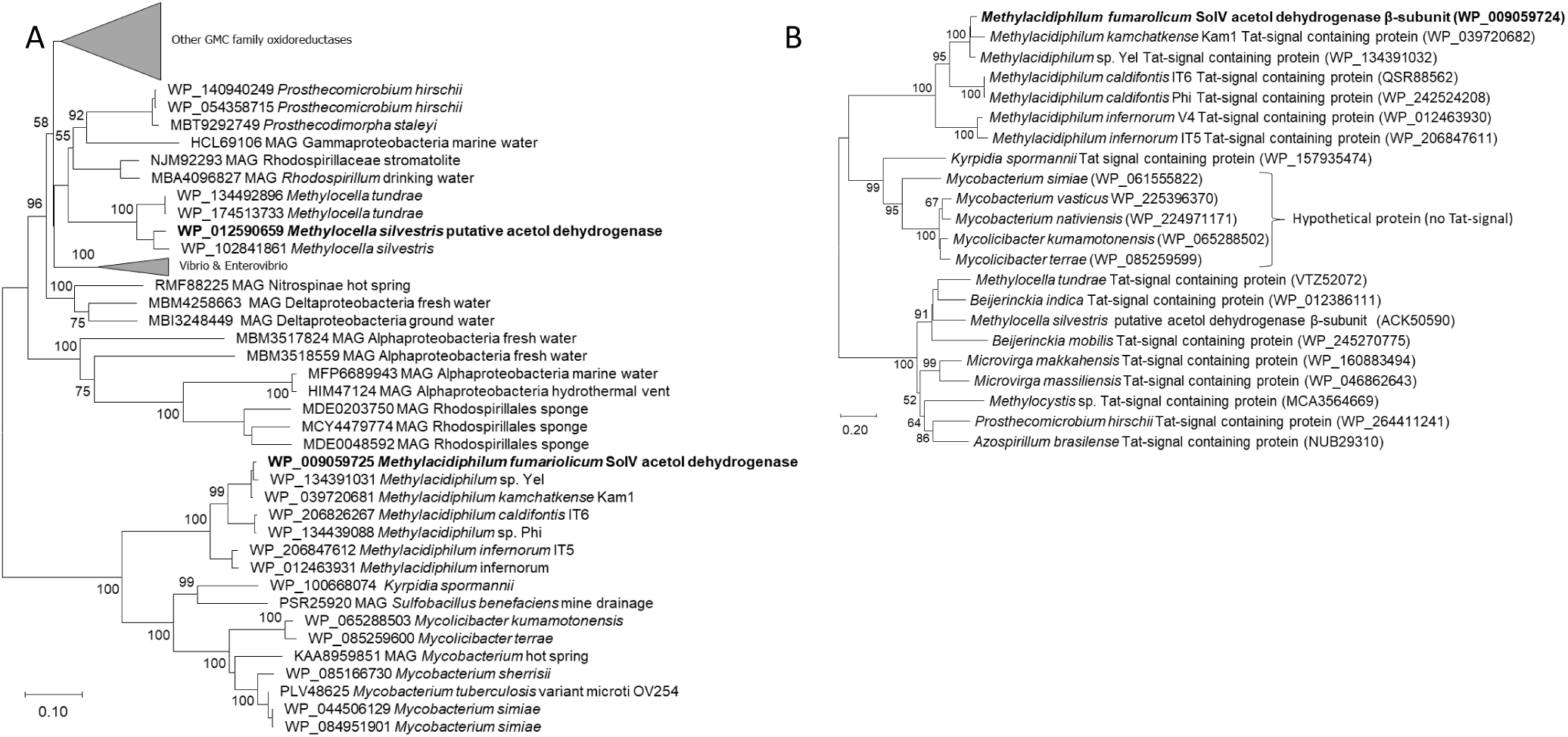
**A.** Phylogenetic tree of the new acetol dehydrogenase with other members of the GMC family oxidoreductases. All sequences show the FAD and FeS cluster binding motifs. The evolutionary history was inferred using the Neighbor-Joining method. The optimal tree with the sum of branch length = 8.74814709 is shown. The percentage of replicate trees in which the associated taxa clustered together in the bootstrap test (500 replicates) is shown next to the branches. The tree is drawn to scale, with branch lengths in the same units as those of the evolutionary distances used to infer the phylogenetic tree. The evolutionary distances were computed using the JTT matrix-based method and are in the units of the number of amino acid substitutions per site. The analysis involved 101 amino acid sequences. All positions with less than 45% site coverage were eliminated. That is, fewer than 55% alignment gaps, missing data, and ambiguous bases were allowed at any position. There were a total of 522 positions in the final dataset. **B.** Phylogenetic analysis of the novel AceDH β-subunit (bold). The evolutionary history was inferred using the Neighbor-Joining method. The optimal tree with the sum of branch length = 5.385 is shown. The percentage of replicate trees in which the associated taxa clustered together in the bootstrap test (500 replicates) is shown next to the branches. The tree is drawn to scale, with branch lengths in the same units as those of the evolutionary distances used to infer the phylogenetic tree. The evolutionary distances were computed using the Poisson correction method and are in the units of the number of amino acid substitutions per site. The analytical procedure encompassed 22 amino acid sequences. The pairwise deletion option was applied to all ambiguous positions for each sequence pair resulting in a final data set comprising 221 positions.

### Sequence alignments

Clustal-Omega was used to prepare the following multiple sequence alignment of selected GMC family α-subunit sequences. The sequence of the AceDH described in this study (**WP_009059725.1**) is highlighted in bold.

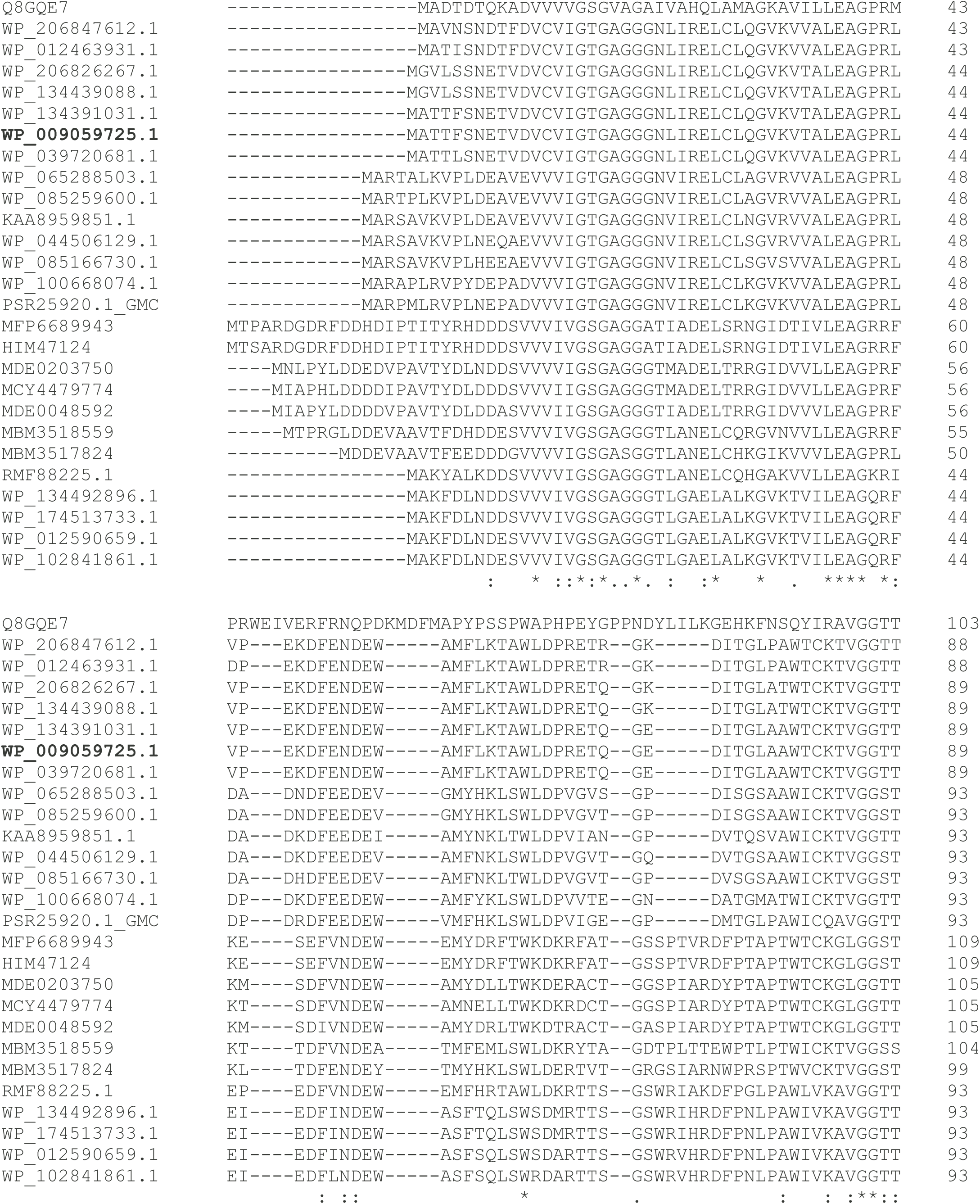
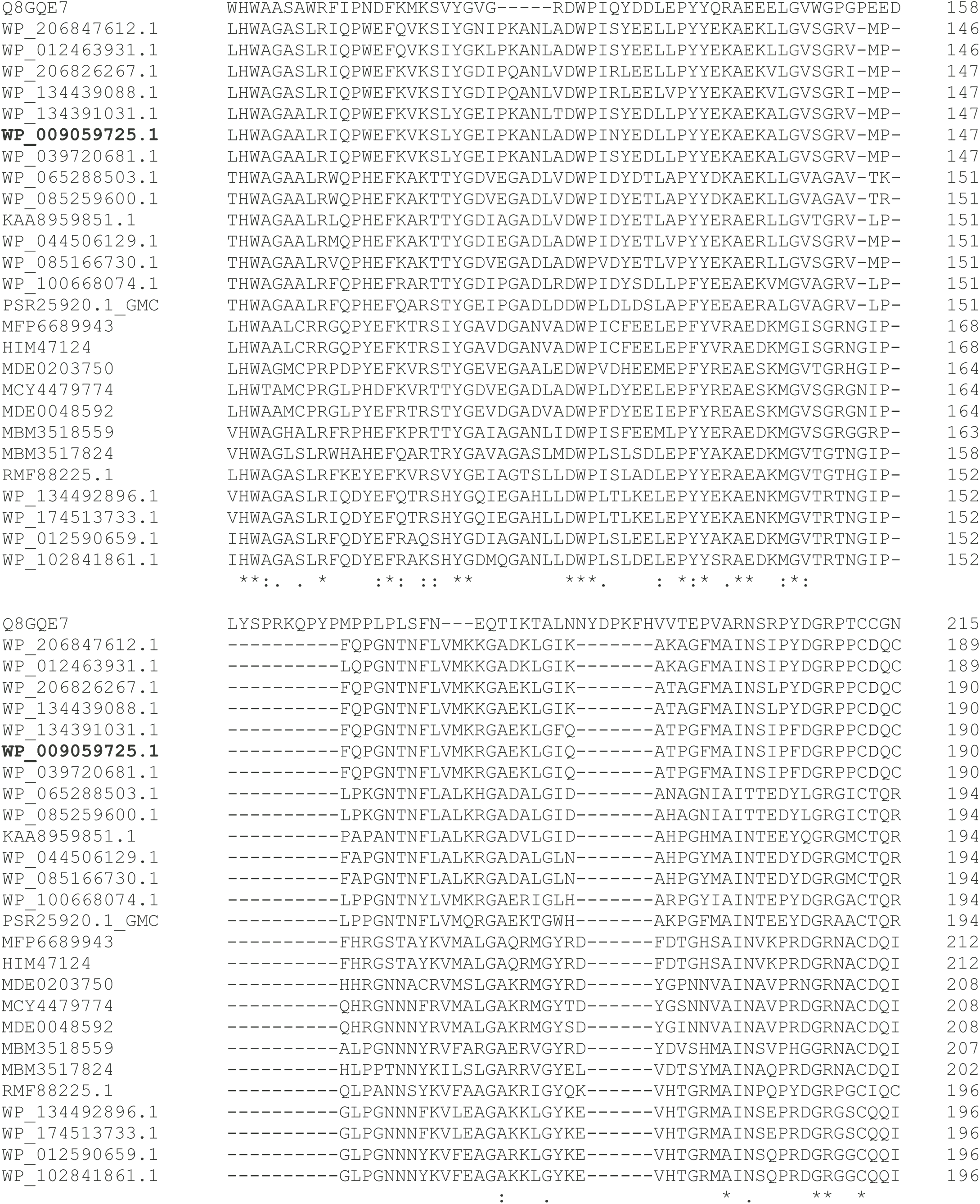
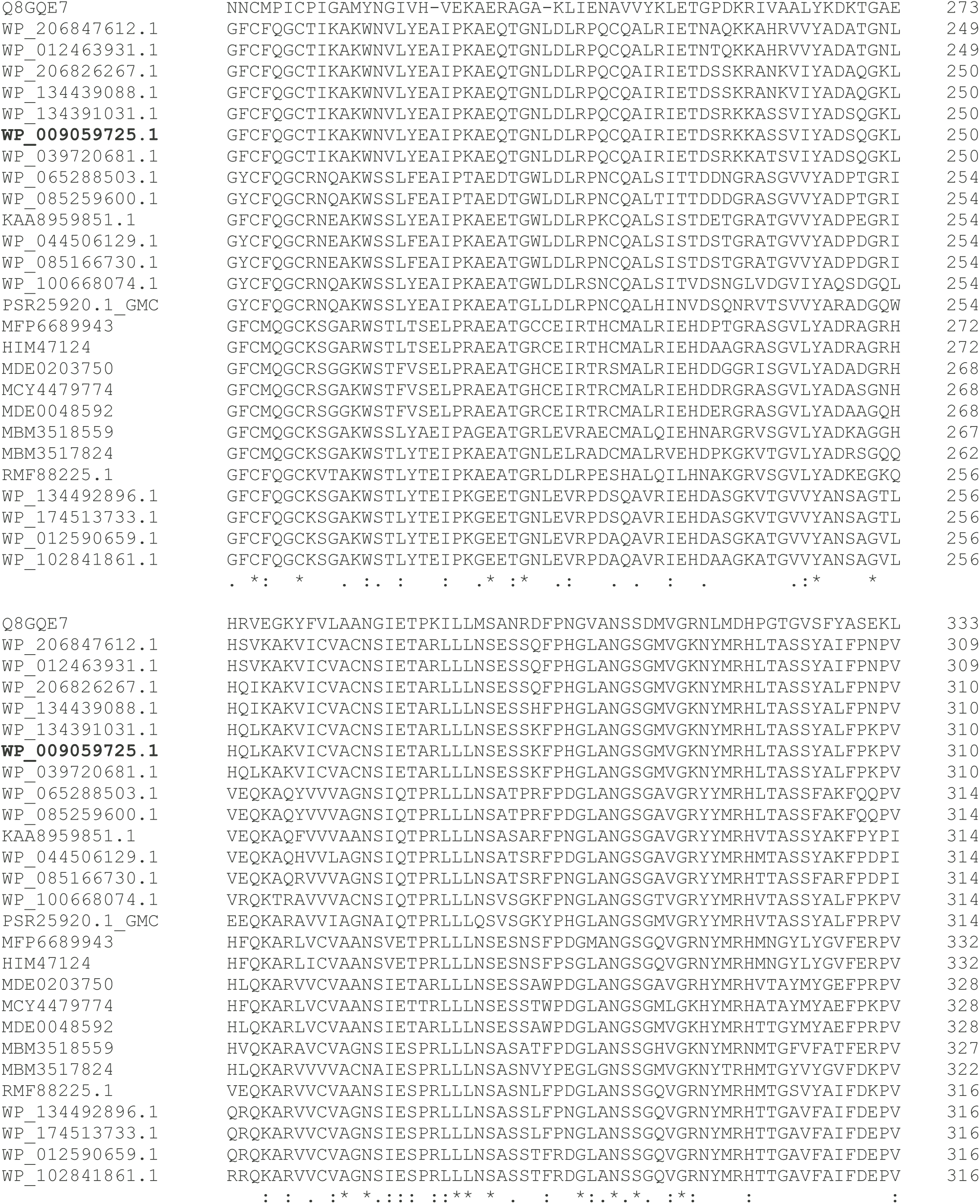
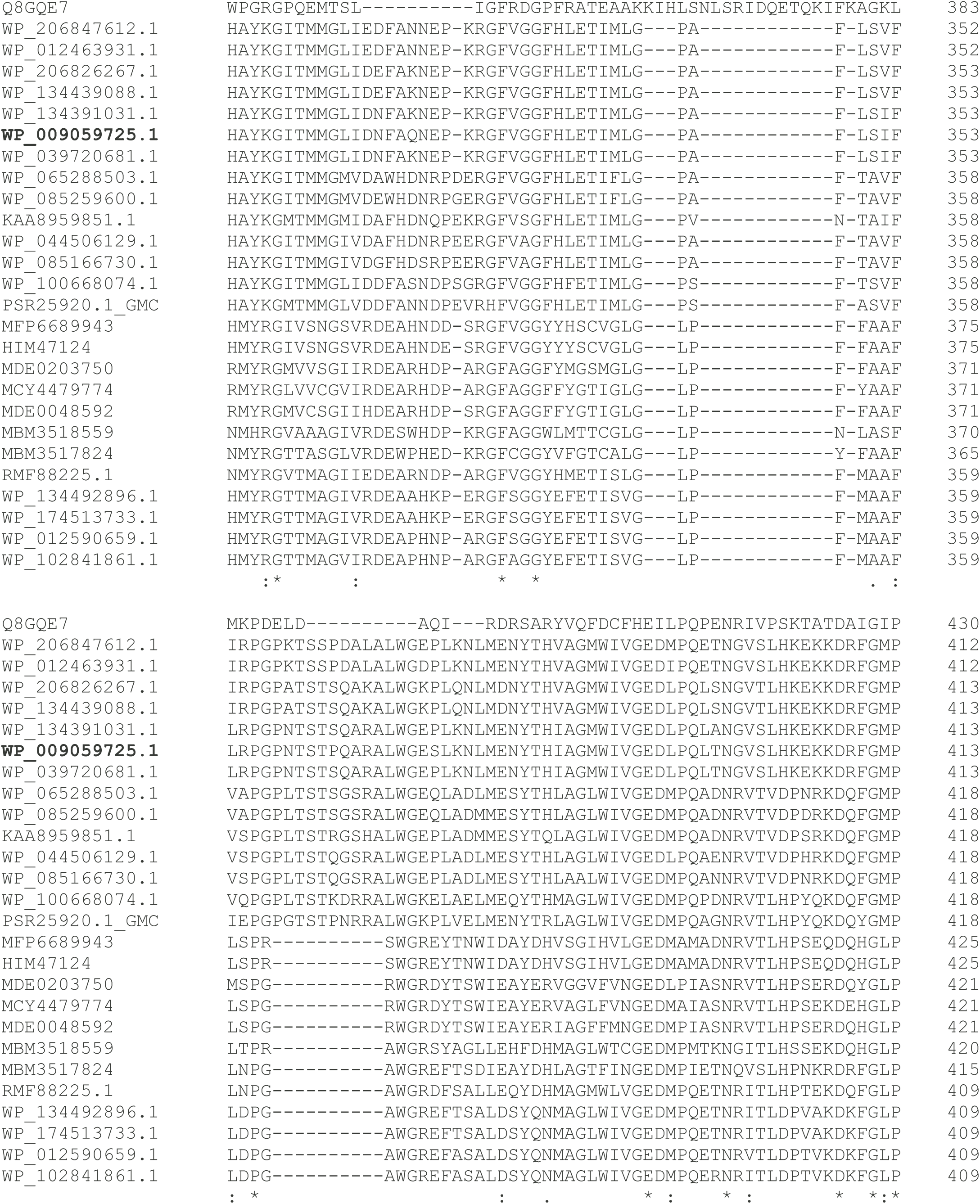
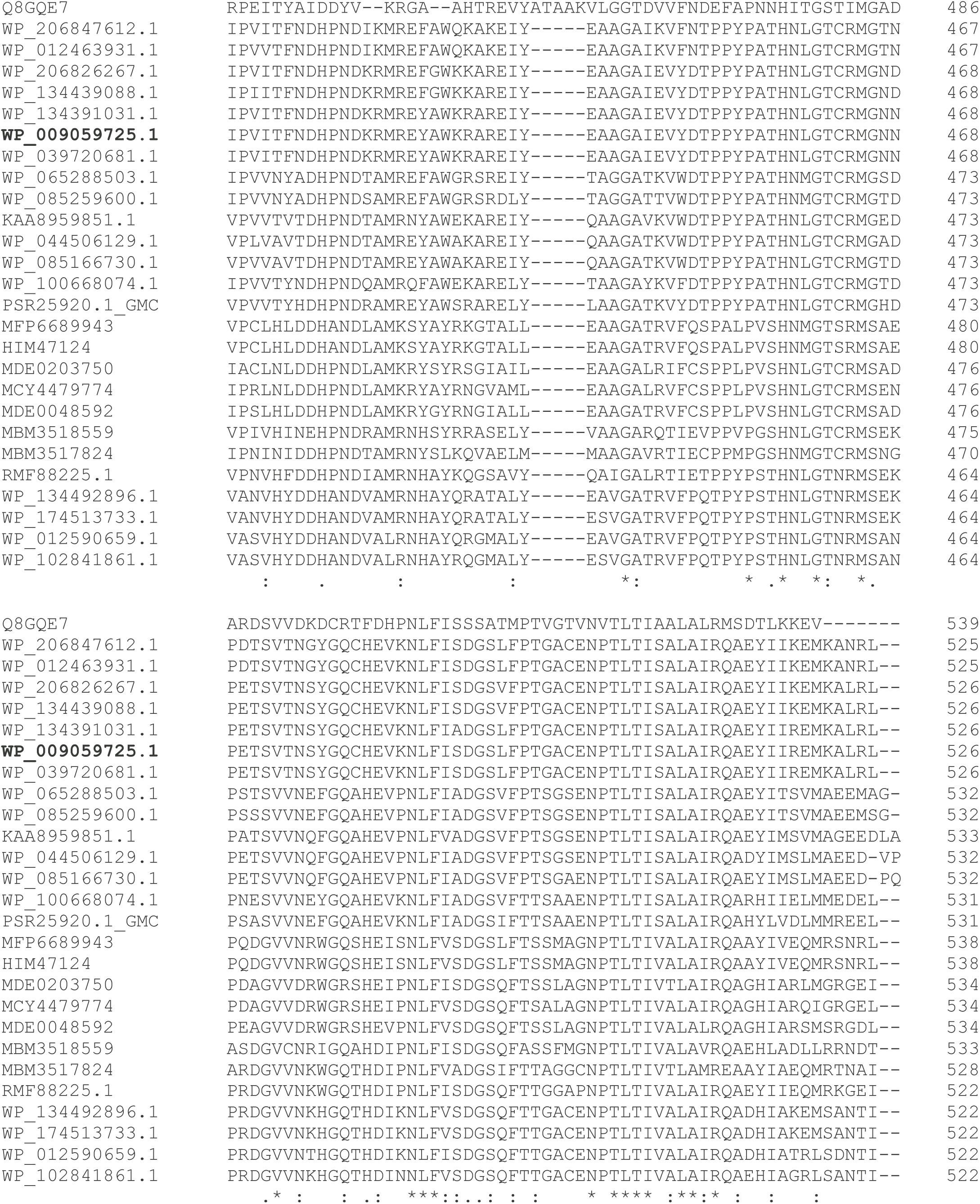
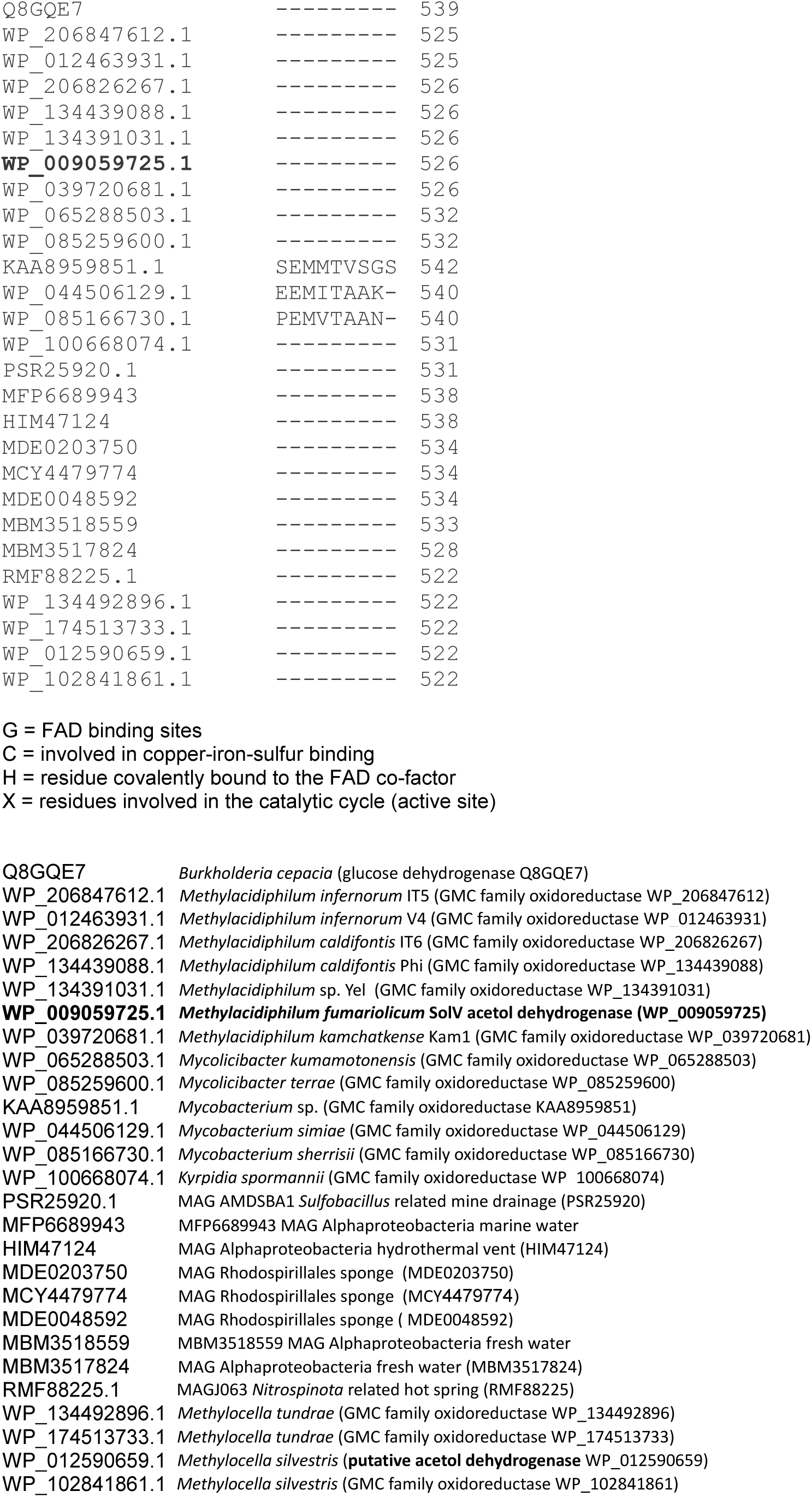

Similarly, the following multiple sequence alignment of the AceDH β-subunit was prepared. Highly conserved C-terminus, including the Methionine residue (Met144 in the crystal structure) involved in binding of the Cu. Putative Tat signal peptide slightly deviating from the [ST]-R-R-x-F-L-K consensus.

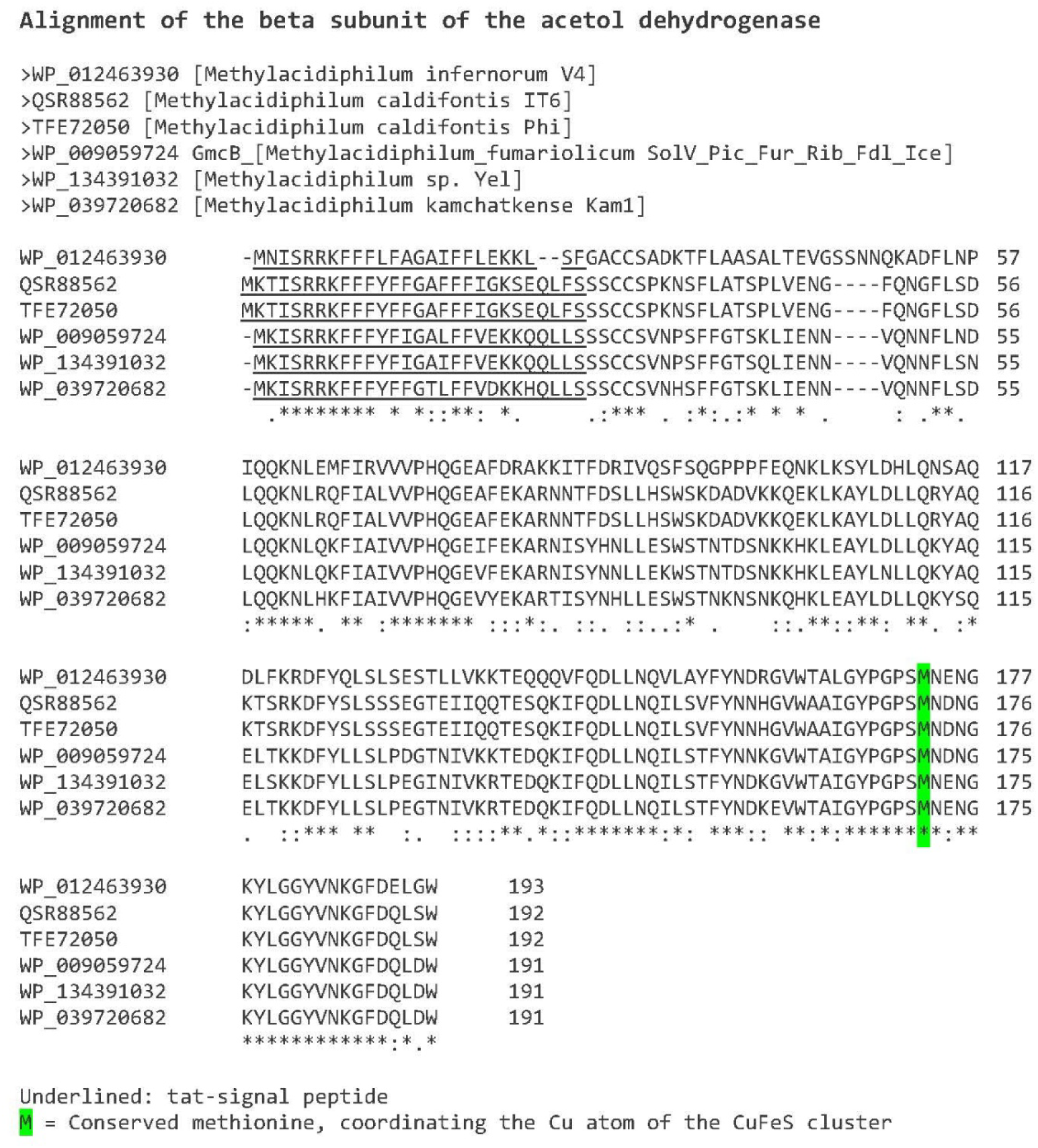

**Extended Data Table 1.** Crystallographic data and model statistics. The outliers include αHis297, which likely has catalytic importance (see Supplemental Information).

|  | AceDH native<br>pdb 9TTU | above Fe edge | above Cu edge |
| --- | --- | --- | --- |
| **Data collection** |  |  |  |
| Space group | $P2_1$ | $P2_1$ | $P2_1$ |
| a, b, c (Å) | 99.6, 92.5, 102.2 | 104.4, 93.7, 107.7 | 104.0, 93.3, 107.5 |
| $\alpha$, $\beta$, $\gamma$ (°) | 90, 116.7, 90 | 90, 116.3, 90 | 90, 116.4, 90 |
| Wavelength (Å) | 0.88560 | 1.72200 | 1.36247 |
| Resolution range (Å) | 65-1.64 (1.67-1.64) | 50-3.70 | 50-3.50 |
| Total no. of reflections | 1,279,261 (30,475) | 102,350 (7,583) | 156,696 (45,378) |
| No. of unique reflections | 185,737 (5,307) | 38,636 (2,835) | 12185 (3363) |
| Completeness (%) | 91.8 (53.0) | 98.7 (99.4) | 99.3 (99.6) |
| Redundancy | 6.9 (5.7) | 2.6 (2.7) | 3.5 (3.6) |
| $\langle I/\sigma(I) \rangle$ | 9.5 (0.5) | 3.5 (1.0) | 5.1 (1.3) |
| $R_{\text{pim}}$ | 0.049 (0.871) | 0.121 (0.889) | 0.155 (0.515) |
| $CC_{1/2}$ | 0.995 (0.356) | 0.973 (0.319) | 0.983 (0.643) |
| **Refinement** |  |  |  |
| Resolution range (Å) | 65-1.8 |  |  |
| Completeness (%) | 98.8 |  |  |
| No. of reflections | 151,401 |  |  |
| Final $R_{\text{cryst}}$ | 0.1749 | | |
| Final $R_{\text{free}}$ | 0.2139 | | |
| No. of non-H atoms |  |  |  |
| Protein | 10,252 |  |  |
| Ligands | 124 (2×[CuFe4S4], 2×FAD) |  |  |
| Solvent | 1038 |  |  |
| R.m.s. deviations |  |  |  |
| Bonds (Å) | 0.008 |  |  |
| Angles (°) | 1.06 |  |  |
| Average $B$ factors (Å<sup>2</sup>) | | | |
| Protein | 32.6 |  |  |
| Ligands | 21.9 |  |  |
| Water | 42.5 |  |  |
| Ramachandran plot |  |  |  |
| Most favoured (%) | 97.2 |  |  |
| Allowed (%) | 2.6 |  |  |
| Outliers (%)* | 0.2 |  |  |
| Rotamer outliers (%) | 0.4 |  |  |

**Extended Data Table 2.** EPR and EXAFS fit results.

| Spin Hamiltonians for EPR simulations |  |  |  |  |  |  |  |  |
| --- | --- | --- | --- | --- | --- | --- | --- | --- |
| | Sim. | $\mathbf{g} = [g_1, g_2, g_3]$ | $\mathbf{gStrain} = [\sigma_1, \sigma_2, \sigma_3]$ | Nucleus 1 | $A_1$ (MHz) | Nucleus 2 | $A_1$ (MHz) | %Weight |
| **X-band** | Cu,N | 2.0273, 1.9841, 1.9672 | 0.036, 0.116, 0.0175 | $^{63,65}\text{Cu}^a$ | 22.7 | $^{14}\text{N}$ | 15.7 | 100 |
| | Cu, H | 2.0273, 1.9841, 1.9672 | 0.051, 0.116, 0.0175 | $^{63,65}\text{Cu}$ | 22.7 | $^1\text{H}$ | 21.7 | 100 |
| | 1 Cu | 2.0273, 1.9841, 1.9672 | 0.051, 0.116, 0.0175 | $^{63,65}\text{Cu}$ | 22.7 | – | – | 100 |
| | 2 Cu | a: 2.0240, 1.9841, 1.9672<br>b: 2.0289, 1.9841, 1.9672 | 0.051, 0.116, 0.0175<br>0.051, 0.116, 0.0175 | $^{63,65}\text{Cu}$<br>$^{63,65}\text{Cu}$ | 23.9<br>24.9 | –<br>– | –<br>– | 42%<br>58% |
| **Q-band** | Cu, H | 2.0266, 1.9799, 1.9672 | 0.030, 0.124, 0.163 | $^{63,65}\text{Cu}$ | 22.7 | $^1\text{H}$ | 21.7 | 100 |
| a) Hyperfine coupling reported for the $^{63}\text{Cu}$ isotope only. Hyperfine coupling of $^{65}\text{Cu}$ is found from the nuclear gyromagnetic ratio $\gamma(g_{\text{H}}(^{65}\text{Cu})/g_{\text{H}}(^{63}\text{Cu})) = 1.0711$. | | | | | | | | |

| Fe EXAFS fitting, $E_0 = 7117.7$ eV | | | | | |
| --- | --- | --- | --- | --- | --- |
| | Path | $N$ | $\sigma^2$ ($\text{\AA}$) | $R$ ($\text{\AA}$) | $R$-factor |
| As isolated | Fe – S4 | $3.5 \pm 0.3$ | $0.003 \pm 0.001$ | $2.28 \pm 0.08$ | 0.004 |
| | Fe – Fe3 (short) | 0.7 | $0.003 \pm 0.001$ | $2.53 \pm 0.03$ | |
| | Fe – Fe1 (long) | 2.6 | $0.003 \pm 0.001$ | $2.72 \pm 0.01$ | |
| Acetol adsorbed | Fe – S4 | $3.5 \pm 0.4$ | $0.003 \pm 0.001$ | $2.28 \pm 0.01$ | 0.005 |
| | Fe – Fe3 (short) | 0.7 | $0.004 \pm 0.001$ | $2.50 \pm 0.05$ | |
| | Fe – Fe1 (long) | 2.6 | $0.004 \pm 0.001$ | $2.72 \pm 0.01$ | |
| Dithionite Reduced | Fe – S4 | $3.3 \pm 0.3$ | $0.003 \pm 0.001$ | $2.28 \pm 0.01$ | 0.003 |
| | Fe – Fe3 (short) | 0.8 | $0.003 \pm 0.001$ | $2.53 \pm 0.02$ | |
| | Fe – Fe1 (long) | 2.5 | $0.003 \pm 0.001$ | $2.72 \pm 0.01$ | |

| Cu EXAFS fitting, $E_0 = 8983.0$ eV | | | | | |
| --- | --- | --- | --- | --- | --- |
| | Path | $N$ | $\sigma^2$ ($\text{\AA}$) | $R$ ($\text{\AA}$) | $R$-factor |
| As-isolated | Cu - S3 | $2.5 \pm 0.4$ | $0.002 \pm 0.001$ | $2.27 \pm 0.01$ | 0.007 |
| | Cu - Fe | 1.0 | $0.002 \pm 0.001$ | $2.59 \pm 0.01$ | |
| Acetol adsorbed | Cu - S3 | $3.0 \pm 0.5$ | $0.005 \pm 0.002$ | $2.28 \pm 0.02$ | 0.006 |
| | Cu - Fe | 1.2 | $0.004 \pm 0.002$ | $2.57 \pm 0.02$ | |
| Dithionite Reduced | Cu - S3 | $3.1 \pm 0.5$ | $0.006 \pm 0.002$ | $2.28 \pm 0.02$ | 0.005 |
| | Cu - Fe | 1.7 | $0.007 \pm 0.002$ | $2.57 \pm 0.02$ | |

$S_0^2$ was fixed as 0.9. $\Delta E_0$ was refined as a global fit parameter, returning a value of $4.0 \pm 1.0$ eV. The $k$ range is 2–11 $\text{\AA}^{-1}$ ( $\Delta R \approx 0.17$ $\text{\AA}$ ) for each sample. The coordination number $N$ of Cu-Fe is constrained as 1 is determined from the crystallographic structure for the as-isolated sample.

**Extended Data Table 3.**
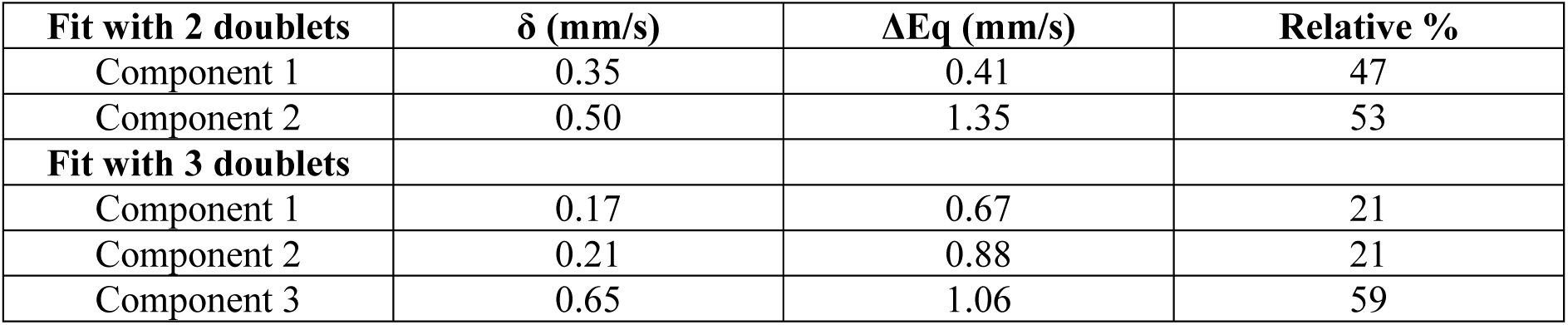
Representative Mössbauer fits.

| Fit with 2 doublets | $\delta$ (mm/s) | $\Delta E_Q$ (mm/s) | Relative % |
| --- | --- | --- | --- |
| Component 1 | 0.35 | 0.41 | 47 |
| Component 2 | 0.50 | 1.35 | 53 |
| Fit with 3 doublets |  |  |  |
| Component 1 | 0.17 | 0.67 | 21 |
| Component 2 | 0.21 | 0.88 | 21 |
| Component 3 | 0.65 | 1.06 | 59 |

**Extended Data Table 4.** GC-MS detection of methylglyoxal product and controls.

| Sample ID | Enzyme<br>(4.44 μM) | PES (2 mM) | Acetol (4.48 μM) | Methylglyoxal<br>(30 μM) | PFBOA<br>(5 mg) | Methylglyoxal<br>detected |
| --- | --- | --- | --- | --- | --- | --- |
| LD-1412-A | ✓ | ✓ | ✓ | ✗ | ✓ |  |
| LD-1512-A | ✗ | ✓ | ✓ | ✗ | ✓ | No |
| LD-1512-B | ✗ | ✓ | ✗ | ✓ | ✓ |  |
| LD-1512-C | ✗ | ✓ | ✗ | ✗ | ✓ | No |
| LD-1512-D | ✗ | ✓ | ✓ | ✗ | ✗ | No |
| LD-1512-E | ✗ | ✓ | ✗ | ✓ | ✗ | No |

